# Changes in SWEET-mediated sugar partitioning affect photosynthesis performance and plant response to drought

**DOI:** 10.1101/2024.04.12.589172

**Authors:** Emilie Aubry, Gilles Clément, Elodie Gilbault, Sylvie Dinant, Rozenn Le Hir

**Author notes:** **Corresponding author:** Dr R. Le Hir. Co-authors email addresses: Emilie Aubry, Gilles Clément, Elodie Gilbault, Sylvie Dinant.

## Abstract

Sugars, produced through photosynthesis, are at the core of all organic compounds synthesized and used for plant growth and response to environmental changes. Their production, transport, and utilization are highly regulated and integrated throughout the plant life cycle. The maintenance of sugar partitioning between the different subcellular compartments and between cells is important in adjusting the photosynthesis performance and response to abiotic constraints. Here, we investigated, in Arabidopsis, the consequences of the disruption of four genes coding for SWEET sugar transporters (SWEET11, SWEET12, SWEET16, and SWEET17) on plant photosynthesis and response to drought. Our results show that mutations in both *SWEET11* and *SWEET12* genes lead to an increase of cytosolic sugars in mesophyll cells and phloem parenchyma cells, which impacts several photosynthesis-related parameters. Further, our results suggest that in the *swt11swt12* double mutant, the sucrose-induced feedback mechanism on stomatal closure is poorly efficient. On the other hand, changes in fructose partitioning in mesophyll and vascular cells, measured in the *swt16swt17* double mutant, positively impact gas exchanges, probably through an increased starch synthesis together with higher vacuolar sugar storage. Finally, we propose that the impaired sugar partitioning, rather than the total amount of sugar observed in the quadruple mutant, is responsible for the enhanced sensitivity upon drought. This work highlights the importance of considering SWEET-mediated sugar partitioning rather than global sugar content in photosynthesis performance and plant response to drought. Such knowledge will pave the way to design new strategies to maintain plant productivity in a challenging environment.

## Introduction

Sugars are the cornerstone of plant growth, development, and response to environmental changes. They are produced by photosynthesis and are then used as carbon skeletons for several anabolic reactions of both primary (e.g. starch and cellulose synthesis) and specialized metabolisms (Louveau and Osbourn 2019). In addition, they act as primary signals in transduction pathways and can serve as osmoprotectants and ROS scavengers in response to abiotic constraints (Saddhe et al. 2021). To achieve this wide range of functions, sugars should be transported from their synthesis site (i.e., photosynthetic source leaves) to their site of use (i.e., sink organs) through the combined action of plasmodesmata and a complex set of active and passive transporters such as sugar will eventually be exported transporters (SWEET) and sucrose transporters (SUC). At all times of the day, plants are constantly adjusting their production, transport, and utilization (e.g., energy production, storage) of sugars to sustain appropriate growth. Indeed, if sink organs are not active enough or sugar loading in source leaves is impaired, sugars accumulate in source leaves, and photosynthesis is inhibited. This phenomenon has been described as the metabolic feedback regulation of photosynthesis (Paul and Pellny 2003). On the other hand, an increased demand from sink organs can enhance photosynthesis (Ainsworth and Bush 2011). At the cell level, manipulations of the vacuole-cytosol sugar exchanges also modify the plant photosynthesis performance in a similar manner. For instance, photosynthesis is promoted in the absence of *PtaSUT4* coding for a tonoplastic sugar exporter or when *AtTST1*, responsible for loading sugars inside the vacuole, is overexpressed (Wingenter et al. 2010; Frost et al. 2012). It has been suggested that a decrease in the cytosolic sugar content is taking place in both cases. At the opposite, the suppression of the expression of *StTST3.1* a tonoplastic sugar importer, potentially leading to an increased cytosolic sugar content, negatively affects leaf photosynthesis in potato (Liu et al. 2023a). In the same line, the overexpression of the chloroplastic glucose facilitator pGlcT2 or the concurrent loss-of-function of pGlcT1 and maltose exporter MEX1, in which the cytosolic glucose availability is proposed to be increased, displays impaired photosynthetic properties (Cho et al. 2011; Valifard et al. 2023).

Due to the global climate change that the earth is facing, recurring or prolonged periods of drought are taking place and negatively affecting plant growth, yield, health and survival (Ghadirnezhad Shiade et al. 2023). When plants face drought stress, several parallel mechanisms take place in the source leaves. The leaves first close their stomata to reduce transpiration. This lowers the CO_2_ absorption and, therefore, lowers photosynthesis, contributing to increased reactive oxygen species (ROS) production. To compensate for the photosynthesis decrease, the source leaves break their starch reserves to produce soluble sugars that will be transported to sink organs such as roots, in which they will be used for growth, and they will be stored in the vacuole thereby maintaining osmotic potential and turgidity (Kaur et al. 2021). The soluble sugars produced will also be used as ROS scavengers to alleviate their negative effects (Keunen et al. 2013). Therefore, the transport of sugars between organs and subcellular compartments constitutes an important control point to ensure an appropriate answer for the plant to drought. Despite the high number of sugar transporters identified in plants, there are relatively few reports about their contribution to the plant response to drought stress (Kaur et al. 2021). Especially it has been shown that the expression of members of the sucrose transporter (SUC/SUT) family could be up or downregulated in response to drought, depending on the phloem loading strategy (apoplasmic versus symplasmic loaders) (Xu et al. 2018). More precisely, in Arabidopsis, the expression of *SUC2*, coding for a sucrose transporter localized at the plasma membrane and responsible for phloem loading, and of *SUC4*, coding for a tonoplastic sucrose transporter, are upregulated in response to drought (Durand et al. 2016; Xu et al. 2018). In addition, the expression of several genes coding for members of the SWEET transporter family, namely *SWEET11*, *SWEET12*, *SWEET13,* and *SWEET15*, is increasing in Arabidopsis leaves in response to water deficit (Durand et al. 2016). These results suggest that improved phloem loading and vacuolar sugar storage are required for plant response to water deprivation. It has been shown that several members of the early response to dehydration six-like (ERDL6) transporter family, which export sugar from the vacuole lumen into the cytosol (Poschet et al. 2011; Khan et al. 2023), are also upregulated in response to water deprivation (Yamada et al. 2010; Slawinski et al. 2021). This supports the important role of vacuole-to-cytosol sugar exchanges in the plant response to drought. In the same line, we recently showed that both in below- and aboveground organs, the carbohydrate distribution mediated by the fructose-specific SWEET17 transporter is crucial for the plant response to drought (Valifard et al. 2021, 2024). Logically, the *swt17* mutant displays a reduced tolerance to drought than the wild type when grown in soil (60% or 40% field capacity) (Valifard et al. 2021). On the other hand, the heterologous overexpression of apple *MdSWEET17* in tomato (*Solanum lycopersicon*) enhances drought tolerance by water withholding for 15 days via fructose accumulation (Lu et al. 2019). Finally, it was recently established that the phosphorylation of SWEET11 and SWEET12 by the SnRK2 kinase in leaves is required to regulate the plant root-to-shoot ratio under drought stress by withholding water for 21-25 days, thereby promoting sucrose transport towards the roots (Chen et al. 2022). As a consequence, the double mutant *swt11swt12* has been shown to be less resistant to drought (Chen et al. 2022).

These examples illustrate that intercellular and intracellular sugar transport should be highly coordinated to achieve proper photosynthesis in normal growth conditions and maintain growth in response to drought. However, previous reports only addressed the effects of a disruption of plasma membrane sugar transporters or tonoplastic sugar transporters. In this work, we propose to go a step further by exploring the simultaneous disruption of SWEET11 and SWEET12 transporters, which are involved in phloem loading (Chen et al. 2012), and of the tonoplastic SWEET16 and SWEET17 transporters controlling the cytosolic soluble sugar availability, on the plant photosynthetic performance and response to drought stress.

## Materials and Methods

### Plant material and growth conditions

The following Arabidopsis lines have been used in this work: Columbia-0 accession (used as wild-type plants and hereafter referred as WT), the *sweet11-1sweet12-1* and the *sweet16-4sweet17-1* double mutants (hereafter referred as *swt11swt12* and *swt16swt17*, respectively) (Le Hir et al. 2015; Aubry et al. 2022) and the *sweet11-1sweet12-1sweet16-4sweet17-1* quadruple mutant (hereafter referred as *swt-q*) (Hoffmann et al. 2022). Seeds were stratified at 4°C for 72 hours in 0.1 % agar solution to synchronize germination. Plants were grown in a growth chamber in short days (8 hours day/16 hours night and 150 µmol m^-2^ s^-1^ brought by fluorescent tubes) at 21/19°C (day/night temperature) with 65% relative humidity, for the thermal and PAM imaging, the gas exchanges measurements and the starch quantification. Alternatively, for the metabolomic analysis (under normal and drought conditions), the targeted gene expression analysis, histochemical analysis, and fluorometric quantification of GUS activity, seeds were sown on peat moss soil plugs in the *Phenoscope* phenotyping robot (https://phenoscope.versailles.inrae.fr/) under short days (8 hours day/16 hours night and 230 µmol m^-2^ s^-1^) at 21/17°C (day/night temperature) with 65% relative humidity as described in Tisné et al. (2013) under different water treatment. More precisely, seeds were let grown for 8 days with a soil water content (SWC) of 100 %. Height days after sowing (DAS) they were transferred to the *Phenoscope.* Watering was maintained through individual pot weight at a specific soil water content (SWC) as described in Tisné et al. (2013). Three watering conditions were tested: ‘60% SWC’ corresponds to 60% of the maximum SWC (representing 4.1g H_2_O per g of dry soil) as a well-watered control condition (stably reached and maintained from 12DAS), ‘30% SWC’ (i.e. 30% of the maximum SWC: 1.6g H2O per g of dry soil) corresponding to a mild drought stress (stably reached and maintained from 16 DAS) and ‘25% SWC’ (i.e. 25% of the maximum SWC: 1.1g H_2_O per g of dry soil) corresponding to a medium intensity drought stress (stably reached and maintained from 17 DAS). Plants were grown in these conditions until 31 DAS and entire rosettes were harvested at 31 DAS for the subsequent experiments. Thanks to daily pictures, the *Phenoscope* also gives access to cumulative growth parameters, such as Projected Rosette Area (PRA), as well as descriptive traits, such as the rosette-encompassing convex hull (Convex Hull Area). Both parameters allow for the calculation of the rosette compactness (PRA/Convex Hull Area), which reflects the rosette morphology (Marchadier et al. 2019).

### Pulse amplitude modulation (PAM) and thermal imaging

The chlorophyll fluorescence was measured using the IMAGING-PAM Maxi version (Heinz Walz GmbH, Effeltrich, Bayern, Germany) driven by the ImaginWin version 2.41a software and equipped with the measuring head consisted of LED-Array illumination unit IMAG-MAX/L and a CCD camera IMAG-MAX/K4 mounted on IMAG-MAX-GS stand. The plants were kept in the dark for a minimum of 20 min prior to the measurements. The rapid light curves were performed by exposing the plants to different light intensities from 1 PAR (µmol photons m^-2^ s^-1^) to 701 PAR according to the manufacturer’s default protocol. The complete rosette was included in the area of interest (AOI), and from the measured values, the derived Fv/Fm ratio, the effective quantum yield of PSII [Y(II)], the electron transport rate (ETR) and the quantum yield of non-photochemical quenching [Y(NPQ)] were calculated according to the equipment manual and represent the mean values of all pixels within an AOI.

Thermal images were obtained using a FLIR A320 infrared camera (Inframetrics, FLIR Systems, North Billerica, MA, USA) equipped with a 16 mm lens. Temperature resolution was below 0.1 °C at room temperature. Straight temperature images generated by the camera software were used based on manufacturer calibration. Leaf emissivity was set to 0.970. For observations, the camera was mounted vertically at approximately 30 cm above the leaf canopy. It was connected to a laptop equipped with the ThermaCAM Researcher Pro 2.9 software for picture acquisition and analysis. For each plant, the total rosette was analyzed, and the minimum (Min), maximum (Max), and the difference maximum-minimum (Max-Min) temperature were given by the software.

### Gas-exchange measurements

A LI-COR 6800F (LI-COR, Lincoln, NE, USA) infrared gas analyzer, equipped with a light source, was used for measuring the gas exchanges. Measurements were performed from 9:00 to 12:00. Leaves were clamped into a 2 cm² chamber. The irradiance level of light was set at 150 µmol photons m^-2^ s^-1^. CO_2_ concentration was set at 400 ppm at constant leaf temperature (around 27°C) and constant vapor pressure deficit (VPD) (around 1.5 kPa). The net CO_2_ assimilation (A_n_), the stomatal conductance (g_sw_), the intercellular CO_2_ concentration (C_i_), and the transpiration rate (E) were measured. The values of the water use efficiency (WUE) and intrinsic water use efficiency (WUEi) were calculated as the A_n_/E and A_n_/g_sw_ ratio, respectively.

### Starch quantification

Starch was quantified from approximately 15-19 mg of whole rosette leaves grown for 43 days in short days and previously lyophilized. The samples were extracted by using a hydroalcoholic extraction consisting of a first step of Ethanol 80% at 90°C for 15 min, a second step of Ethanol 30% at 90°C for 15 min, and a last step of water at 90°C for 15 min. Between each extraction round, a centrifugation step at 5 000 rpm for 3 minutes allowed the collection of the supernatants that could be used for soluble sugars quantification. The pellet was then dried in an oven at 50°C for 3h. After hydrolyzing the starch pellet using thermostable α-amylase and amyloglucosidase, glucose was quantified by enzymatic reaction as described in Bergmeyer and Bernt (1974).

### Histochemical analysis of GUS activity

The lines expressing pSWEET11:SWEET11-GUS or pSWEET12:SWEET12-GUS (Chen et al. 2012) and pSWEET16:SWEET16-GUS or pSWEET17:SWEET17-GUS (Guo et al. 2014) in Col-0 background were used to assess SWEET11, SWEET12, SWEET16 and SWEET17 expression pattern in response to different watering regimes (i.e., 60% SWC, 30 % SWC and 25 % SWC). The histochemical GUS staining was performed according to Sorin et al. (2005). Pictures were taken using an Axio zoom V16 microscope equipped with a Plan-NEOFLUARZ 2.3x/0.57 FWD 10.6mm objective and the ZEN (blue edition) software package (Zeiss, https://www.zeiss.com/).

### Metabolomic analysis

All steps were adapted from the original protocols detailed in Fiehn (2006, 2016) and Fiehn et al. (2008). The detailed protocol is available in Amiour et al. (2012). Briefly, 30 mg of ground frozen rosette leaves per genotype and condition were used for metabolome analysis by an Agilent 7890A gas chromatograph (GC) coupled to an Agilent 5977B mass spectrometer (MS). The column was an Rxi-5SilMS from Restek (30 m with 10 m Integra-Guard column). During the extraction phase, 4µg/ml of ribitol were added to each sample to allow metabolite absolute quantification. Then, a response coefficient was determined for 4 ng each of a set of 103 metabolites, respectively, to the same amount of ribitol. This factor was used to estimate the absolute concentration of the metabolite in what we may call a “one-point calibration.” Standards were injected at the beginning and end of the analysis. Data were analyzed using AMDIS (http://chemdata.nist.gov/mass-spc/amdis/) and TargetLynx softwares (Waters Corp., Milford, MA, USA). For each genotype and condition, between 7 to 8 plants were analyzed. The table containing the raw data of the metabolomic profiling is available under the link https://doi.org/10.57745/VD55XO.

### RNA isolation and cDNA synthesis

RNAs were prepared from whole rosette leaves subjected to different water regimes and grown in short-day conditions as described above. Samples were frozen in liquid nitrogen before being grounded with a mortar and pestle. Powders were stored at -80°C until use. Total RNA was extracted from frozen tissue using TRIzol reagent (Thermo Fisher Scientific, 15595-026, https://www.thermofisher.com) and treated with DNase I, RNase-free (Thermo Fisher Scientific, EN0521, https://www.thermofisher.com). cDNA was synthesized by reverse transcribing 1 µg of total RNA using RevertAid H minus reverse transcriptase (Thermo Fisher Scientific, EP0452, https://www.thermofisher.com) with 1 µl of oligo(dT)18 primer (100 pmoles) according to the manufacturer’s instructions. The reaction was stopped by incubation at 70 °C for 10 min.

### RT-qPCR experiment

Transcript levels were assessed for four independent biological replicates in assays with triplicate reaction mixtures by using specific primers either designed with the Primer3 software (http://bioinfo.ut.ee/primer3-0.4.0/primer3/) or taken from the literature (Supplementary Table S1). qPCR reactions were performed in a 96-well transparent plate on a Bio-Rad CFX96 Real-Time PCR machine (Bio-Rad) in 10 µl mixtures each containing 5 µl of Takyon™ ROX SYBR^®^ MasterMix dTTP Blue (Eurogentec, UF-RSMT-B0710, https://www.eurogentec.com/), 0.3 µl forward and reverse primer (30 µM each), 2.2 µl sterile water and 2.5 µl of a 1/30 dilution of cDNA. The following qPCR program was applied: initial denaturation at 95°C for 5 min, followed by thirty-nine cycles of 95°C for 10 sec, 60°C for 20 sec, 72°C for 30 sec. Melting curves were derived after each amplification by increasing the temperature in 0.5°C increments from 65°C to 95°C. The Cq values for each sample were acquired using the Bio-Rad CFX Manager 3.0 software package. The specificity of amplification was assessed for each gene, using dissociation curve analysis, by the precision of a unique dissociation peak. If one of the Cq values differed from the other two replicates by > 0.5, it was removed from the analysis. The amplification efficiencies of each primer pair were calculated from a 10-fold serial dilution series of cDNA (Supplementary Table S1). Four genes were tested as potential reference genes: *APT1* (At1g27450), *TIP41* (At4g34270), *EF1α* (At5g60390), and *UBQ5* (At3g62250). The Normfinder algorithm (Andersen et al. 2004) was used to determine the gene most stably expressed among the different genotypes analyzed, namely *EF1α* in this study. The relative expression level for each genotype was calculated according to the ΔCt method using the following formula: average E_t_^-Cq(of^ ^target^ ^gene^ ^in^ ^A)^/E_r_^-Cq(of reference^ ^gene^ ^in^ ^A)^, where E_t_ is the amplification efficiency of the target gene primers, E_r_ is the reference gene primer efficiency, A represents one of the genotypes analyzed.

### Statistical analysis

The statistical analyses were performed using either a student t-test or a two-way ANOVA combined with Tukey’s comparison post-test using R software. A *p*-value of < 0.05 was considered as significant. For the metabolome analysis, the Rflomics R package coupled with a shiny application and developed by INRAE was used and is available in Github (https://github.com/RFLOMICS/RFLOMICS). Briefly, the raw data were transformed using square root, and a differential expression analysis was performed using the lmfit model of the limma package (Ritchie et al. 2015). The Rflomics package was also used to re-analyze the RNA-seq results obtained by Khan et al. (2023). Filtering and normalization strategies were performed as described in Lambert et al. (2020).

## Results

### Photosynthetic performance is impaired in *swt11swt12* double mutant

It has been proposed that an excess sugars can trigger negative feedback regulation of leaf photosynthesis (Paul and Foyer 2001; Ainsworth and Bush 2011). In agreement with this, the disruption or overexpression of genes coding for sugar transporters, which potentially impact the cytosolic sugar availability, induces modifications of the photosynthetic performance (Wingenter et al. 2010; Frost et al. 2012; Khan et al. 2023; Liu et al. 2023b; Valifard et al. 2023). In an effort to gain more knowledge on the physiological consequences of impaired SWEET-mediated sugar transport, we measured growth and photosynthesis performance in the *swt11swt12*, *swt16swt17,* and *swt-q* mutants and in the wild-type Col-0 (WT). The growth of the different mutant lines was first estimated by measuring the projected rosette area (PRA) after 43 days of growth under short-day (SD) conditions. In these conditions, both *swt11swt12* and *swt-q* mutants is smaller than that of the WT, while no significant changes of the PRA of *swt16swt17* mutant are measured (Supplementary Fig. S1). Several parameters related to photosynthesis performance and nonphotochemical quenching using PAM imaging (rapid light curve) were measured. While the *swt16swt17* mutant did not show any significant difference compared to the WT plants, we show that, along with an increase in the light intensity, the effective quantum yield of PSII photochemistry (Y(II)) is significantly lower in *swt11swt12* and *swt-q* mutants compared to WT plants (Fig. 1A). This is accompanied by a significant decrease of the electron transport rate (ETR) of both mutants compared to the wild type (Fig. 1B). Moreover, we measured the fraction of energy dissipated in form of heat via the regulated non-photochemical quenching mechanisms [Y(NPQ)]. In wild-type plants, this fraction is logically increasing together with the increase of the light intensity (Fig. 1C). In both *swt11swt12* and *swt-q* mutants, the Y(NPQ) values are globally significantly higher compared to WT, showing that more energy is dissipated in the form of heat in the mutant lines. Thus, avoiding PSII damage that could have occurred because of the impaired mutant capacity to efficiently use light. Altogether, these results point to the fact that impairment of both *SWEET11* and *SWEET12* expression leads to impaired photosynthetic efficiency.

**Fig. 1.**
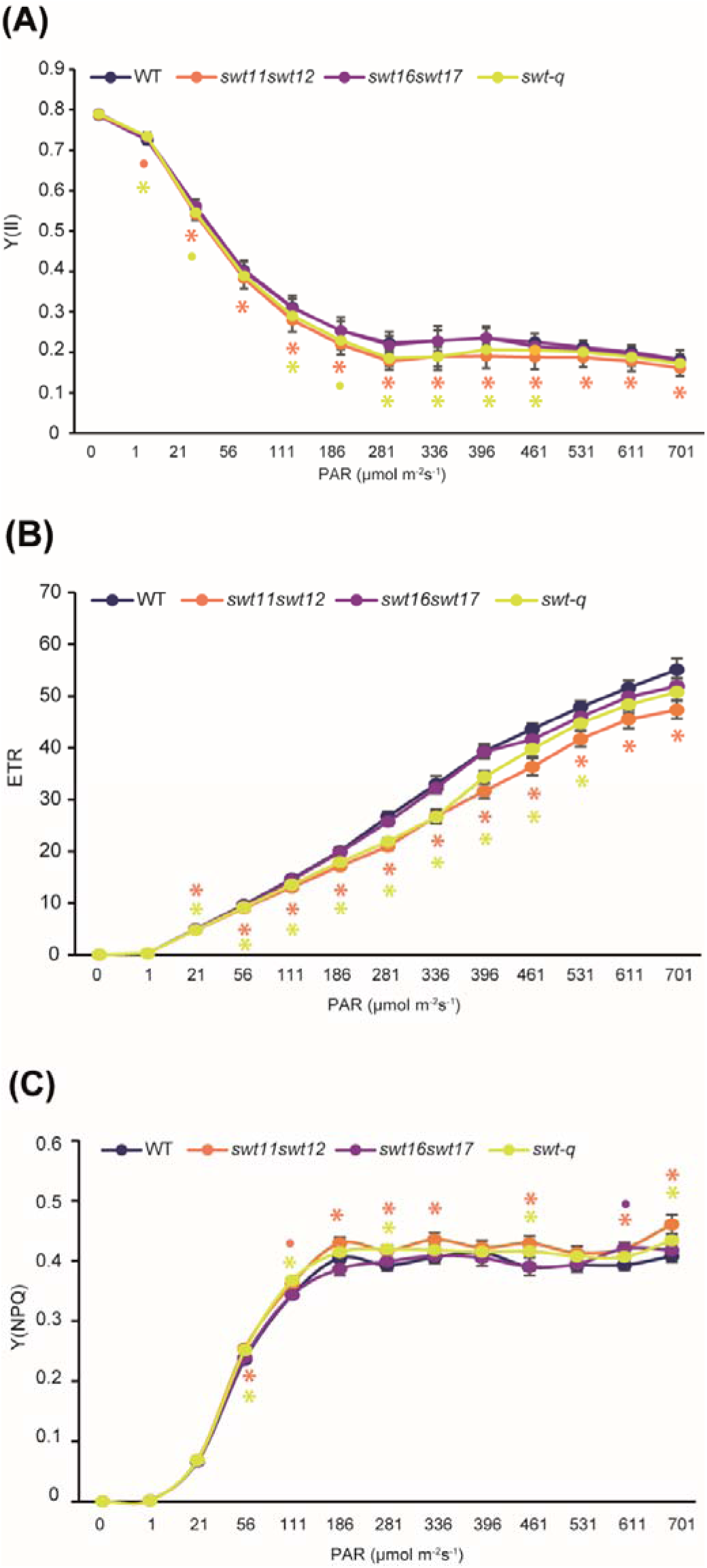
Disruption of *SWEET11* and *SWEET12* leads to a loss of photosynthesis efficiency. The effective quantum yield of PSII [Y(II)] (A), the electron transport rate (ETR) (B), and the quantum yield of non-photochemical quenching [Y(NPQ)] (C) were determined using a light curve of increasing PAR intensity performed on wild type (WT), *swt11swt12*, *swt16swt17* and *swt-q* mutants. Values are means ± SE (n = 12 plants). Significant differences were calculated using a student’s *t*-test with * *p*<0.05 and • 0.05 < *p* < 0.09.

### Gas-exchange parameters are altered in *sweet* mutant lines

Since feedback between CO_2_ assimilation and photosynthetic activity has been shown, we further explored if this lower photosynthetic capacity could be linked to a limitation in the CO_2_ assimilation. At the leaf level, the gas exchanges in the different mutants were assessed (Fig. 2). Under ambient light conditions, the stomatal conductance (g_sw_) along with the transpiration rate (E) of the *swt11swt12* and *swt-q* mutants tend to increase compared to WT plants, albeit not significant (Fig. 2A-B). In addition, a tendency, albeit not significant, for a lower net CO_2_ assimilation (A_net_) was measured in both *swt11swt12* and *swt-q* lines (Fig. 2C). On the contrary, the assimilation rate as well as the stomatal conductance and the transpiration rate of the *swt16swt17* double mutant are significantly improved compared to the WT (Fig. 2A-C). Further, we show that the intracellular CO_2_ concentration (Ci) is significantly higher in all the *sweet* mutant lines compared to wild-type plants (Fig. 2D). This increase is logical since Ci calculation is more dependent on variations in g_sw_ and E than that of A_net_, according to the equation of von Caemmerer and Farquhar (1981). Therefore, the improved A_net_ observed in *swt16swt17* is not important enough to induce a decrease in Ci values in this genotype. Nonetheless, this higher Ci suggests that more CO_2_ substrate is available for assimilation in the *sweet* mutant lines. From these parameters, the water-use efficiency (WUE), which represents the ratio of biomass produced per unit of water consumed (A/E ratio), and the intrinsic water-use efficiency (WUEi), which represents the ratio of net assimilation over the stomatal conductance (A/g_sw_ ratio), were calculated (Fig. 2E and F). The *swt11swt12* mutant displays a reduced WUE and WUEi compared to the wild type, suggesting that less biomass is produced per unit of water loss (Fig. 2E and F). This is consistent with the smaller projected rosette area measured for this genotype (Supplementary Fig. S1). On the other hand, the *swt16swt17* and the *swt-q* mutants did not display any significant changes in WUE, while the WUEi is significantly decreased compared to wild-type plants (Fig. 2E and F). These results suggest that the stomata functioning and/or development could be affected in these lines. By using thermal imaging, we were able to show that the maximum rosette temperature is significantly reduced in the *swt11swt12* double mutant, and consequently, the difference between the maximum and the minimum temperature is also significantly reduced in this line (Supplementary Fig. S2B-C). On the other hand, no significant changes in these parameters are observed in *swt16swt17* and *swt-q* mutants (Supplementary Fig. S2A-C). Altogether, these results show that, in normal growth conditions, the impairment of the SWEET-mediated sugar transport impacts the equilibrium between CO_2_ assimilation and transpiration.

**Fig. 2.**
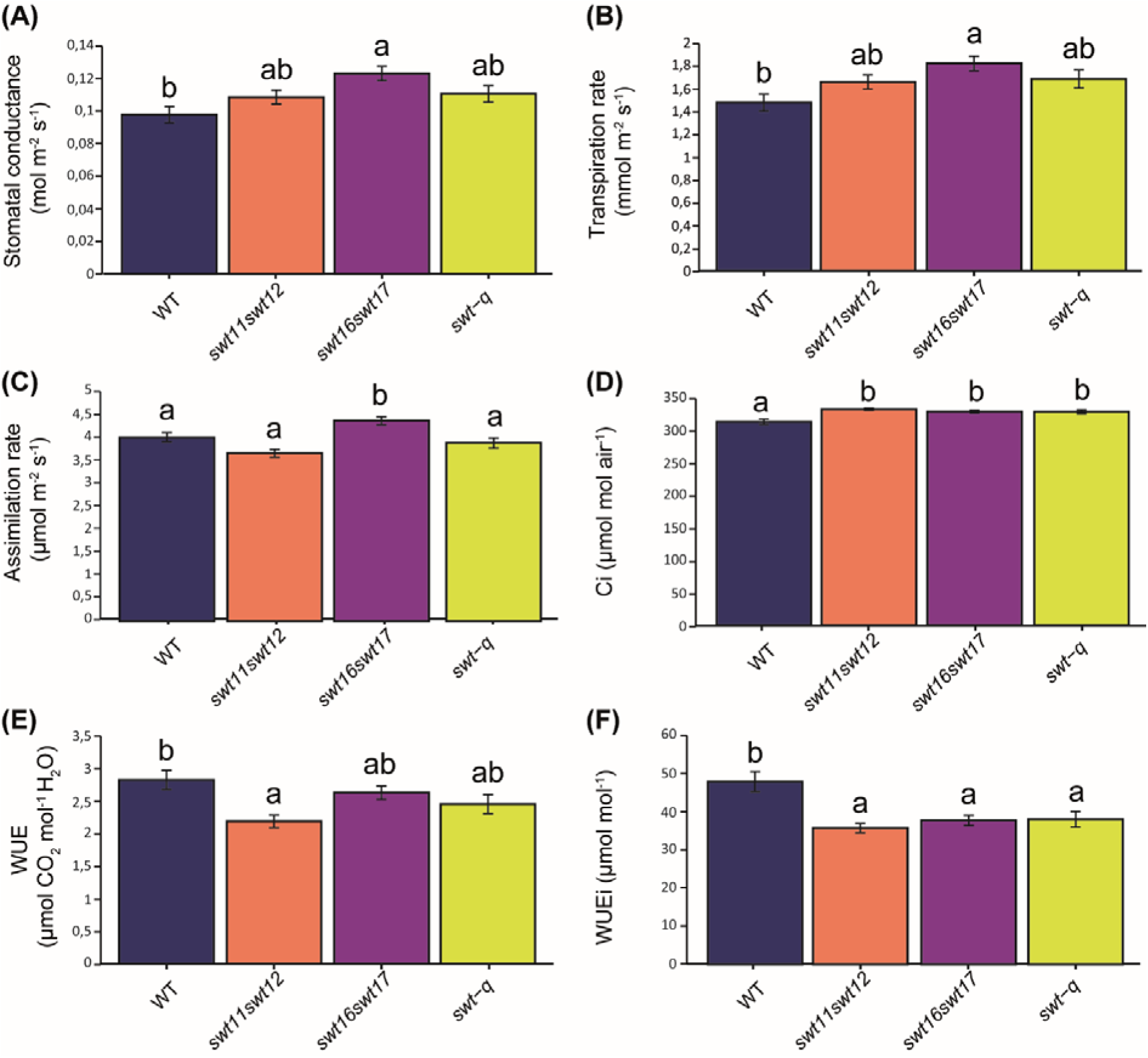
Assimilation and transpiration rates are impaired in *sweet* mutants. Barplots showing the stomatal conductance (A), the transpiration rate (B), the assimilation rate (C), the intracellular CO_2_ concentration (C_i_) (D), the ratio assimilation/transpiration or water use efficiency (WUE) (E), and assimilation/stomatal conductance or intrinsic water use efficiency (WUE_i_) (F) of wild type (WT), *swt11swt12*, *swt16swt17* and *swt-q* mutants grown in short-day photoperiod. Means ± SE are shown (n = 12 plants from two independent cultures; 3 leaves per plant were measured). A one-way ANOVA combined with the Tukey’s comparison post hoc test was performed. The values marked with the same letter were not significantly different, whereas different letters indicate significant differences (*p* < 0.05).

#### SWEET transporters are differently expressed in rosette leaves

We exploited the published single-cell transcriptomics dataset of short day-grown leaves (Kim et al. 2021) to access the cell-specific expression of *SWEET11*, *SWEET12*, *SWEET16* and *SWEET17* (Supplementary Fig. S3). Among the four genes tested, we could see that *SWEET11* is the most expressed in the rosette leaves grown in short days, followed by *SWEET17* (Supplementary Fig. S3A). On the other hand, *SWEET12* and *SWEET16* are lowly expressed (Supplementary Fig. S3A). More precisely, *SWEET11* is mainly expressed in a cluster of cells related to phloem parenchyma/procambium (cluster 10) and phloem parenchyma/xylem (cluster 18) (Supplementary Fig. S3B). At the same time, *SWEET12* is co-expressed with *SWEET11* in the phloem parenchyma/xylem cells (cluster 18) (Supplementary Fig. S3C). Surprisingly, *SWEET16* is almost exclusively expressed in the cluster 13 related to epidermal cells (Supplementary Fig. S3D) while an expression of *SWEET17* is found in nearly all the different leaves cell types including the cluster 16 related to guard cells (Supplementary Fig. S3E). We completed these data by performing a histochemical analysis of GUS activity on lines expressing *SWEET-GUS* fusion proteins driven by their native promoter (Supplementary Fig. S3F-I). As expected from the single-cell transcriptomic analysis, *SWEET11* was expressed mainly in the leaf vascular system (Supplementary Fig. S3F). However, no expression of either *SWEET12* or *SWEET16* was observed in our growing conditions (Supplementary Fig. S3G and H). The absence of expression of these proteins, while a low expression is measured at the transcript level, suggests that both genes could be down-tuned at the translational or post-translational level in our growing conditions. Finally, *SWEET17* expression was mainly observed in the vascular system of the leaves as well as in cells surrounding the vasculature (Supplementary Fig. S3I).

### Sugars and organic acids of the TCA cycle accumulate in *sweet* mutants

Next, the metabolic status of these lines was assessed by performing a global metabolomic profiling (Fig. 3 and https://doi.org/10.57745/VD55XO) on plants grown in SD photoperiod on the *Phenoscope* to ensure high sample reproducibility. In *swt11swt12* and *swt-q* mutants, the analysis of the global metabolites identified 43 and 47 metabolites significantly different compared to wild-type plants, respectively (Supplementary Table S2). Consistently with previous reports (Chen et al. 2012; Le Hir et al. 2015), a significant accumulation of sucrose (fold-change (FC)=2.12), glucose (FC=1.81), and fructose (FC=1.61) was measured in the *swt11swt12* mutant (Fig. 3A and Supplementary Table S2). A significant increase of the same sugars is also measured in the *swt-q* mutant compared to the WT (FC sucrose =1.87, FC glucose =1.68 and FC fructose =2.75). The content of other sugars, including xylose, myo-inositol, melibiose, raffinose, and mannose was also significantly accumulated in *swt11swt12* and/or *swt-q* mutants, albeit to a lower extent than sucrose, glucose and fructose (1.02<FC<1.20), while the content of stachyose was slightly decreased in these mutants (FC=0.96) (Fig. 3A and Supplementary Table S2). In addition, several organic acids, among which almost all those involved in the TCA cycle (i.e., malate, citrate, succinate and 2-oxoglutarate) were significantly accumulated in both mutant lines (1.08<FC<2.04) (Fig. 3A and supplementary Table S2). On the other hand, the content of several amino acids (e.g., GABA, arginine, leucine, isoleucine, glycine) was slightly decreased only in the *swt-q* mutant line compared to wild-type plants (0.94<FC<0.98) (Fig. 3 and supplementary Table S2). Interestingly, in the *swt16swt17* mutant, the statistical analysis only identified fructose (FC=1.69) as differentially accumulated compared to wild-type plants (Fig. 3A and supplementary Table S2). The content of fructose increased similarly in both double mutants. In contrast, the fructose content doubled in the quadruple mutant compared to the double mutants, suggesting an additive phenotype (Fig. 3B). To complete this metabolomic analysis, an analysis of the starch content of SD-grown leaves was performed and show that the three mutant lines are significantly accumulating starch compared to wild-type plants (Supplementary Fig. S4).

**Fig. 3.**
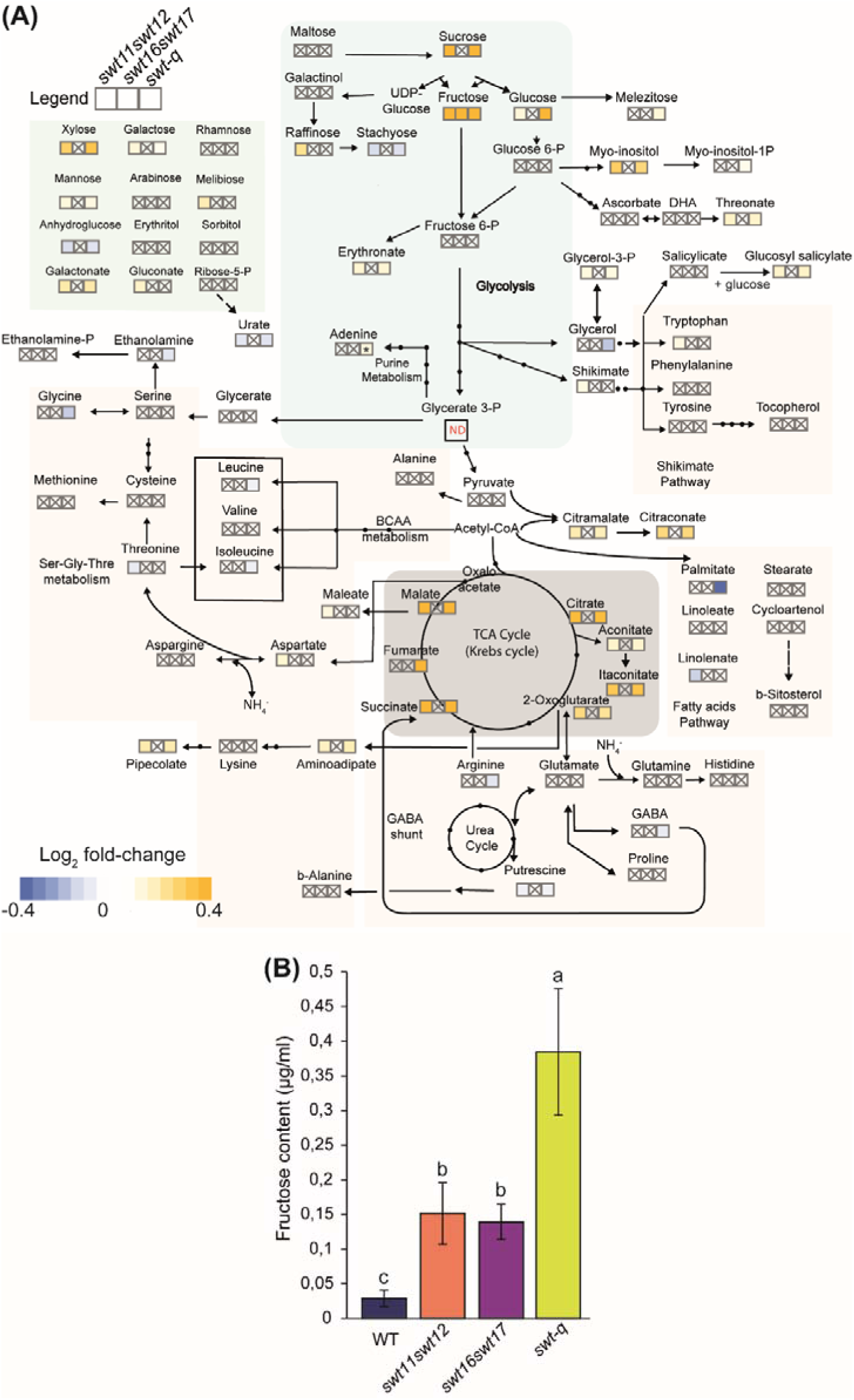
Metabolites involved in sugar metabolism and in the Krebs cycle are accumulating in rosette leaves of *sweet* mutants. (A) Heatmap showing the changes in the metabolite contents in the *swt11swt12*, *swt16swt17,* and *swt-q* mutants grown in the phenotyping robot. Values presented are Log_2_ transformed mutant-to-wild-type ratio (n ≥ 7 per genotype). The color gradient from yellow to blue refers to metabolites accumulated or decreased, respectively, in the mutants compared to the wild type. Only metabolites displaying a statistical difference are shown as Log_2_ fold-change. Values with *p* < 0.05 were considered significantly different. Crossed out boxes: Metabolites showing no statistical differences. ND: not detected. (B) Barplot showing differences in fructose content from the metabolomic analysis in the different mutants. A one-way ANOVA combined with the Tukey’s comparison post hoc test was performed. The values marked with the same letter were not significantly different, whereas different letters indicate significant differences (*p* < 0.05).

### *sweet* mutants exhibit altered expression of genes related to sugar partitioning

As stated above, *swt11swt12 and swt-q* mutants accumulate more soluble sugars and organic acids involved in the TCA cycle, suggesting that glycolysis may also be affected in these mutants. Indeed, the Krebs cycle is fuelled by pyruvate from cytosolic glycolysis performed during the day. On the other hand, the plastidial glycolysis allows the production of energy during the dark period. Thus, we checked, on the same samples as those used for the metabolite profiling, for alterations of the expression of genes coding for the first steps of cytosolic and plastidial glycolysis (Table 1). We also measured the expression of genes involved in tonoplastic and plastidic sugar/triose phosphate transport to get a global picture of the metabolite’s dynamic in the different subcellular compartments (Table 1).

**Table 1.**
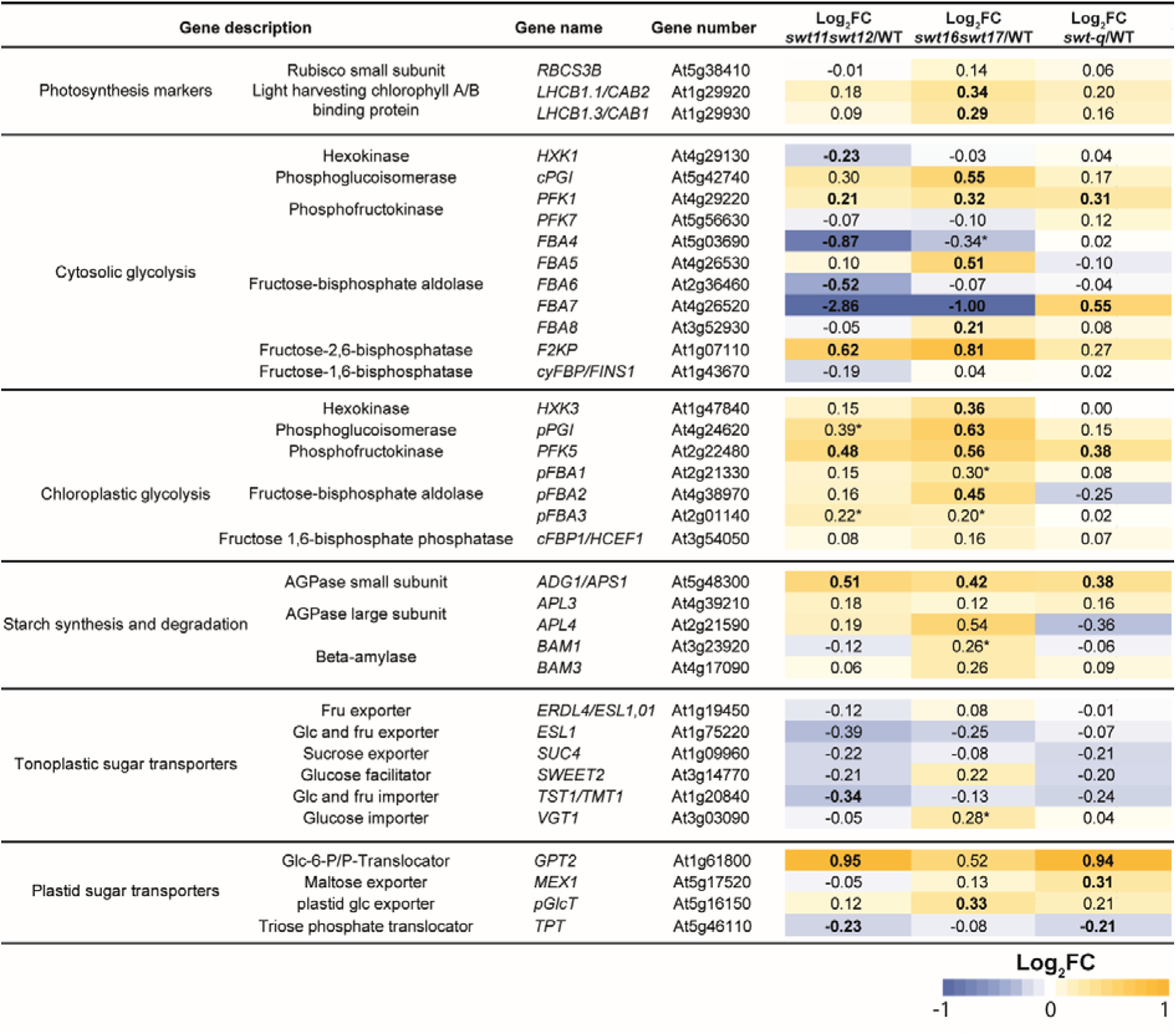
Log_2_-fold changes in the expression of genes involved in sugar homeostasis in the cytosol the chloroplast in *sweet* mutant plants compared to WT grown in short-days photoperiod. Whole rosettes have been harvested and used for mRNA extraction and subsequent qPCR experiments. Bold-scripted Log_2_FC indicate significant fold changes (*p* < 0.05), while stars indicate Log2FC values for which p is comprised between 0.05 and 0.09 according to a double-sided t-test.

During the day, sucrose synthesis in the cytosol in leaves is fuelled mainly by triose phosphates exported from the chloroplasts by the TRIOSE-PHOSHATE/PHOSPHATE TRANSLOCATOR (TPT) and are first converted into fructose-1,6-biphosphate (Fru-1,6-P_2_) by the FRUCTOSE BIPHOSPHATE ALDOLASE (FBA). Then Fru-1,6-P_2_ is dephosphorylated into fructose-6-phosphate (Fru-6-P) by FRUCTOSE 1,6-BIPHOSPHATASE (cyFBP/FINS1). Conversely, Fru-6-P is phosphorylated into Fru-1,6-P_2_ by PHOSPHOFRUCTOKINASE (PFK). The conversion of Fru1,6-P_2_ into Fru-6-P is under the direct regulation of the fructose 2,6-biphosphate produced by the 6-PHOSPHOFRUCTO-2-KINASE/FRUCTOSE-2,6-BIPHOSPHATASE (F2KP) enzyme (McCormick and Kruger 2015).

In the *swt11swt12* mutant, we measured a significant upregulation of the *F2KP* and *PFK1* expression together with a global significant downregulation of the expression of genes coding for enzymes involved in converting triose phosphate into Fru-1,6-P_2_ (i.e., *FBA4*, *FBA6*, *FBA7*) and for the triose-phosphate translocator (*TPT*) (Table 1). In addition, a significant decrease in the expression of *TONOPLAST SUGAR TRANSPORTER1* (*TST1*) coding for a glucose and fructose importer and an increase in the expression of chloroplastic *GLC6-PHOSPHATE TRANSLOCATOR2* (*GPT2*) are measured (Table 1). Therefore, these results suggest that the phosphorylation/dephosphorylation of the cytosolic fructose-6-P, which is critical for the balance between sucrose and starch production, is impaired in the *swt11swt12* mutant. At the chloroplast level, we could measure a significant upregulation of several genes related to chloroplastic glycolysis (i.e., *pFBA3*, *PFK5*, *pPGI*). In agreement with a previous report showing an accumulation of starch in the *swt11swt12* double mutant grown under SD conditions (Gebauer et al. 2017), a significant upregulation of *ADG1* involved in starch biosynthesis is also shown (Table 1). Interestingly, in the *swt16swt17* mutant, we also measured a significant upregulation of both *F2KP* and *PFK1* expression (Table 1). However, depending on the isoforms, an up- or downregulation of *FBA* genes is observed (Table 1). At the chloroplast level, a significant upregulation of *HEXOKINASE3* (*HXK3*), *PHOSPHOGLUCO ISOMERASE1* (*pPGI*), *PFK5*, *pFBA1*, *pFBA2* and *pFBA3* along with *ADP GLUCOSE PYROPHOSPHORYLASE1* (*ADG1*) and *BETA-AMYLASE1* (*BAM1*), involved in starch biosynthesis and hydrolysis, is measured (Table 1). Finally, a significant upregulation of both *PLASTIDIC GLUCOSE TRANSLOCATOR* (*pGlcT)* coding for a chloroplastic glucose exporter and of *VACUOLAR GLUCOSE TRANSPORTER1* (*VGT1*) coding for a vacuolar glucose importer is observed (Table 1). In the quadruple mutant line, similar to both double mutant lines, a significant upregulation of the cytosolic isoform *PFK1* and the chloroplastic isoform *PFK5* is measured (Table 1). However, unlike both double mutants, the expression of *FBA7* (cytosolic isoform) is also significantly upregulated in the *swt-q* mutant (Table 1). In addition, *ADG1* and *GPT2* are also significantly upregulated in *swt-q* mutant as in *swt11swt12* mutant (Table 1). Finally, an upregulation of *MEX1* coding for a maltose transporter is only measured in the quadruple mutant (Table 1).

Finally, since modifications in gas exchanges were measured in *sweet* mutant lines together with an accumulation of sugars, we also measured the expression of genes related to photosynthesis (i.e., *RBCS3A*, *LCHB1.1*/*CAB2* and *LHCB1.3*/*CAB1*). Interestingly, the sugar-repressed genes *CAB1* and *CAB2* are significantly upregulated in *swt16swt17* mutant compared to wild-type plants (Table 1). However, no change in the expression of photosynthesis-related genes is measured in both *swt11swt12* and *swt-q* mutant lines (Table 1). Altogether these results show that, in addition to modifications of the photosynthesis, the balance between cytosolic and plastidial glycolysis is strongly affected when the expression of *SWEET* transporters is disrupted. The increased expression of *F2KP*, *ADG, PFK5,* and *GPT2* suggests, in particular, that the sugar metabolism is redirected toward starch accumulation.

### The rosette growth of *swt-q* mutants is negatively affected in response to drought stress

Previously, it has been shown that the *swt11swt12* double mutant is more susceptible to drought stress (Valifard et al. 2021; Chen et al. 2022). Consistently, both WUE and WUEi are significantly decreasing in this mutant line (Fig. 2). Alongside, we show that the *swt-q* mutant is also displaying a decreased WUE while accumulating a higher amount of fructose than both double mutant lines (Figs. 2 and 3). Therefore, we checked the *swt-q* mutant response to different drought scenari using the *Phenoscope* phenotyping robot. Three conditions were tested: non-limiting watering conditions (60% SWC), mild stress intensity (30% SWC) and medium stress intensity (25% SWC). The raw data are presented in Supplementary Table S3. When grown at 30% SWC, the projected rosette area (PRA) and the convex hull encompassing the rosette of the wild type decreased compared to a SWC of 60%, albeit not significantly (Fig. 4A-B). A further decrease of the SWC to reach 25% impacts significantly wild-type plants by reducing the PRA and the convex hull area by 35% and 50 %, respectively (Fig. 4A-B). In agreement with previous work (Chen et al. 2022), we show that the *swt11swt12* mutant displays an increased drought sensitivity when comparing the PRA loss between normal watering conditions and mild-intensity stress (30% SWC) (31% for *swt11swt12* versus 11% for wild-type plants) (Supplementary Fig. S5). Moreover, our results show that the PRA loss observed in the *swt16swt17* double mutant is reduced (27%) compared to wild-type plants (42%) between normal watering conditions and 35 %SWC, suggesting a slightly improved resistance to drought (Supplementary Fig. S5). Furthermore, under normal watering conditions, the PRA and the convex hull area of the *swt-q* mutant is significantly smaller than that of the wild type (Fig. 4A-B). When grown under a mild-intensity stress (30 % SWC), the PRA and the convex hull area of the mutant line are more affected than the wild-type plants (Fig. 4A-B) and similar to that of the *swt11swt12* double mutant (Supplementary Fig. S5). When grown under medium-intensity stress (25 % SWC), the PRA and the convex hull area of the *swt-q* mutants are not significantly different from that of the wild-type plants grown in the same condition (Fig. 4A-B). Finally, we show that the PRA loss of WT plants is 35% between both drought stress conditions (30 % SWC and 25 % SWC). In comparison, that of the *swt-q* mutant is reduced by 18% (Fig. 4A). Consistently, we observe for both traits a significant effect of the interaction between genotype and drought stress intensity (GxE) according to the result of a two-way ANOVA (Fig. 4A-B). We calculated the rosette compactness, which reflects the rosette morphology and potential default in leaves elongation, of both genotypes under the different watering conditions (Fig. 4C). Interestingly, the rosette of WT is significantly more compact when plants are subjected to a medium intensity stress (25 % SWC). On the other hand, under normal watering conditions, the rosette of the *swt-q* mutant is smaller but also more compact than that of the wild type (Fig. 4C). However, unlike WT plants, the compactness of the *swt-q* rosette does not significantly change in response to drought stress (Fig. 4C). Overall, these results show that despite having a smaller and more compact rosette in normal watering conditions (60 % SWC) *swt-q* mutants display an increased sensitivity to drought.

**Fig. 4.**
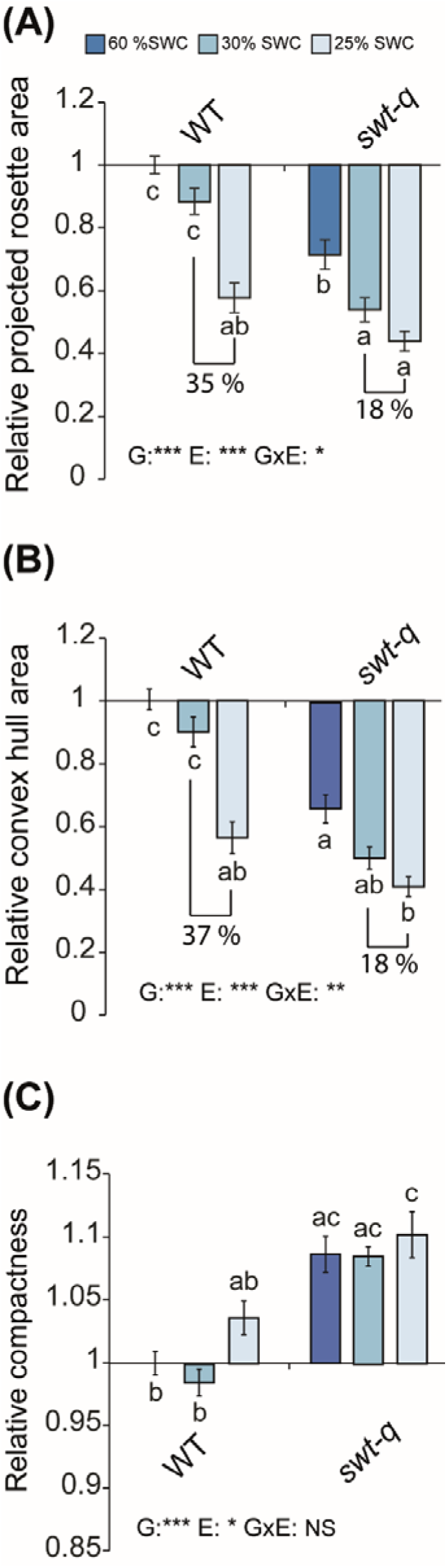
The *swt-q* mutant is more sensitive to drought stress than the wild type. Barplots showing the fold-changes in the projected rosette area (A), the convex hull area (B), and the compactness (C) of the wild type and quadruple mutant grown in short-days photoperiod under different watering regimes: 60% SWC (normal watering conditions), 30 % SWC (mild stress intensity) or 25% SWC (medium stress intensity). The fold changes were determined using the values normalized to the mean of wild-type plants grown at 60% SWC. A two-way ANOVA combined with the Tukey’s comparison post hoc test was performed. The values marked with the same letter were not significantly different, whereas different letters indicate significant differences (*p* < 0.05, *n* ≥ 18 per genotype and condition from 2 independent experiments).

### Metabolic status of *swt-q* mutant and wild-type plants under drought stress

Next, we analyzed the metabolite profiles of wild-type plants and *swt-q* mutant grown under the different watering regimes (Fig. 5 and Supplementary Tables 4-8). As expected, mild and medium-intensity stresses lead to changes in the metabolite content of the rosette of both genotypes (Supplementary Tables 4-7). In WT plants, only 3 metabolites were found to be accumulated in response to mild drought stress compared to non-limiting watering conditions, namely ascorbate (FC=1.65), nicotinate (FC=1.008), and anhydroglucose (FC=1.11) (Fig. 5 and Supplementary Table S4). Ascorbate and nicotinate are involved in oxidative stress and redox reactions, while anhydroglucose results from the hydrolysis of cell wall compounds (Durand et al. 2019). On the other hand, the content of 51 metabolites over a total of 54 was differentially reduced in response to mild intensity stress (30% SWC) (Fig. 5 and Supplementary Table S4), among which 13 amino acids, 10 organic acids as well as sugars (e.g., fructose, trehalose) and phosphorylated sugars (e.g., glucose-6-phosphate, fructose-6-phosphate). In response to medium-intensity drought stress (25% SWC), we identified 52 metabolites differentially altered in comparison with the non-limiting condition (60% SWC), among which 18 metabolites were accumulated, and 34 were reduced (Supplementary Table S4). The content of proline tends to increase in response to medium drought stress, albeit not significantly (Supplementary Fig. S6A). Nonetheless, we found a slight but significant accumulation of the 1-pyrroline-5-carboxylase (precursor of proline) (FC=1.014) along with an increased expression of the *DELTA 1-PYRROLINE-5-CARBOXYLATE SYNTHASE2* (*P5CS2*) gene responsible for the proline synthesis in response to both drought stress conditions (Supplementary Table S5 and Supplementary Fig. S6B). Several organic acids such as ascorbate and intermediate of the Krebs cycle (e.g., citrate, citramalate, fumarate, malate) were also accumulated (Fig. 5 and Supplementary Table S5) (1.02<FC<1.87). Finally, the content of several sugars, already known to be involved in the plant response to drought, was also increased but at a slight level (e.g., raffinose, myo-inositol, galactinol) (1.08<FC<1.19). Similarly, in response to mild drought stress, the content of several amino acids along with sugars and sugar-phosphate was reduced in response to medium drought stress (Fig. 5 and supplementary Table S5). Since the most important effect of drought stress on WT rosette growth was observed in response to 25 % SWC (Fig. 5), we also identified metabolites specifically altered in response to medium drought stress. The content of 15 metabolites were specifically accumulated among which several organic acids (e.g., malate, fumarate, citrate, citramalate, galactonate, erythronate, 1-pyrroline-5-carboxylate) and sugars (e.g., raffinose, myo-inositol, xylitol) (Fig. 5 and Supplementary Fig. S7A). On the other hand, the content of only 5 metabolites were specifically decreasing in response to medium drought stress (i.e., putative hexose-P, C12:0 dodecanoate, galactosylglycerol, U2390.0/04 [1-Benzylglucopyranoside] and a breakdown product of glucosinolate) (Fig. 5 and Supplementary Fig. S7B).

**Fig. 5.**
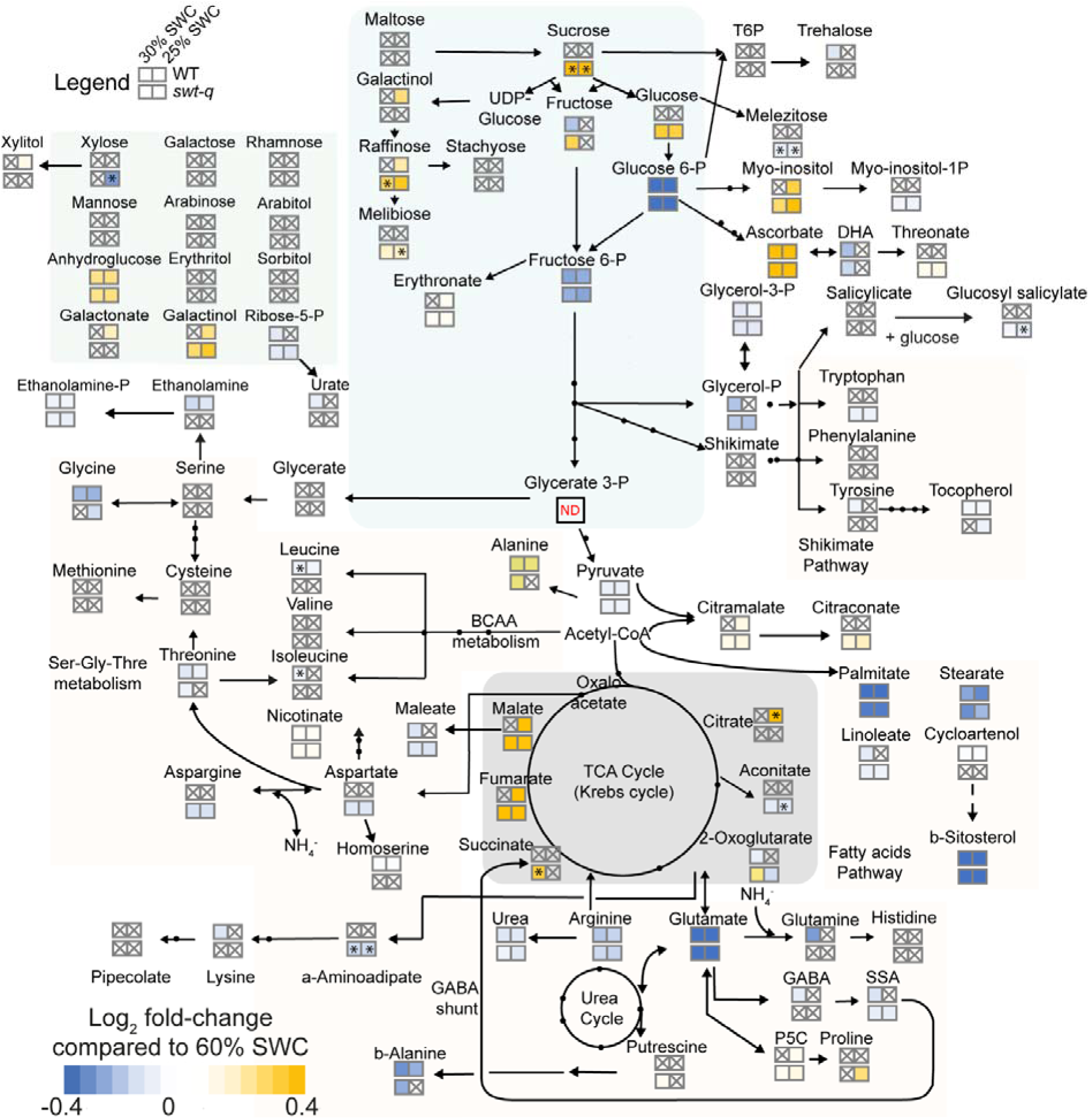
The *swt-q* mutant accumulates more sugars than the wild type in response to drought stress. Heat map of the metabolite content changes in the wild type and the *swt-q* mutant grown in short-day photoperiod under different watering conditions. The orange-to-blue gradient refers to metabolites accumulated (yellow) or decreased (blue) for the wild type and the *swt-q* mutant. The values presented in the two upper squares are Log_2_ fold-change ratio of the wild type grown under 30% SCW (mild stress intensity) (left square) and 25% SWC (medium stress intensity) (right square) compared to the wild type grown under 60 % SWC (normal conditions). The values presented in the two lower squares are Log_2_ fold-change ratio of the *swt-q* plants grown under 30% SCW (mild stress intensity) (left square) and 25% SWC (medium stress intensity) (right square) compared to the *swt-q* grown under 60 % SWC (normal conditions). Additional stars in boxes indicate metabolites significantly different between wild type and *swt-q* mutant. Only metabolites showing a significant difference according to a One-way ANOVA followed by a Tukey HSD post-test are presented (*p* < 0.05, *n* ≥ 7 plants per genotype and condition). Crossed out boxes: metabolites showing no statistical differences. ND: not detected.

In *swt-q* mutants, 22 and 20 metabolites were significantly accumulating when grown under mild drought stress (30 % SWC) or medium drought stress (25 % SWC) compared to non-limiting watering condition, respectively (Fig. 5 and Supplementary Tables S6-7). Nineteen of them were found common between mild and medium intensity drought stress (e.g., raffinose, sucrose, erythronate, 1-pyrroline-5-carboxylate, anhydroglucose, ribonate, malate, citramalate, myo-inositol, fumarate, nicotinate, ascorbate, galactinol and glucopyranose) (Supplementary Tables S6-7). Especially, fumarate was strongly accumulated in both stress conditions (FC=2.91 in 30 % SWC and FC=2.34 in 25 % SWC) (Supplementary Tables S6-7). Only fructose, putrescine and succinate were specifically accumulated in response to mild intensity drought stress (Fig. 5 and Supplementary Table S6-7). Finally, 47 and 45 metabolites were shown to have a decreased content in response to mild and medium drought stress, respectively (Supplementary Tables S6-7). Like for accumulating metabolites, most of the decreasing metabolites were common between both stress conditions (Supplementary Fig. S7 and Supplementary Tables S6-7). Further we show that only proline was specifically accumulating in *swt-q* subjected to 25 % SWC (FC=1.13) (Supplementary Fig. S7C) while xylose, glycine, alpha-tocopherol, galactosylglycerol, a fatty acid methyl ester and an unknown metabolite related to ketohexose were specifically decreasing in response to 25 % SWC (Supplementary Fig. S7D).

Finally, we looked for metabolites responding to the interaction between genotype and environment. Eight metabolites (i.e., isoleucine, leucine, succinate, raffinose and sucrose, a-aminoadipate and melezitose) were shown to be significant to the combination of genotype and environment (60% SWC vs 30 % SWC) (Supplementary Table S8). Especially, unlike wild-type plants, *swt-q* mutants accumulate mainly sucrose in response to medium drought stress (FC=2.38) and a slight accumulation of raffinose (FC=1.15) and an unknown metabolite related to hexose-phosphate (FC=1.008) is also shown (Fig. 5 and Supplementary Table S8). On the other hand, isoleucine and leucine are slightly but significantly accumulating in wild-type plants unlike in *swt-q* mutants (FC=1.02) (Fig. 5 and Supplementary Table S8). When comparing 60 % SWC and 25 % SWC, a specific accumulation of melibiose, sucrose, aconitate, and melezitose is measured in *swt-q* mutants (Fig. 5 and Supplementary Table S8). On the other hand, while citrate significantly accumulates in WT in response to medium drought stress, its content (already high in normal conditions) did not change in *swt-q* mutant (Fig. 5 and Supplementary Table S8).

### In response to drought stress, the genes involved in starch synthesis and degradation are upregulated in *swt-q* mutants compared to WT plants

We further analyzed the swt-q mutant response to drought stress by quantifying the changes of expression of genes related to drought response, photosynthesis/respiration, and those involved in sugar partitioning in the different watering conditions (Table 2 and Supplementary Table S9). In response to drought stress, *swt-q* mutant responds similarly to WT plants by significantly upregulating the expression of *RESPONSE TO DESSICATION 29A* (*RD29a*), *P5CS2,* and *EARLY RESPONSE TO DEHYDRATION SIX-LIKE1* (*ERDL3.07*/*ESL1*), known to be stress markers genes (Table 2). Despite an initial defect in photosynthesis performance (Fig. 1), the photosynthesis marker genes (i.e., *RBSC3B*, *LHCB1.1*/*CAB2* and *LHCB1.3*/*CAB1*) were not significantly regulated in response to drought in the *swt-q* mutant while a slightly significant upregulation of the expression of *RBSC3B* is measured in WT plants in response to mild intensity drought stress (Table 2). Interestingly, a significant increase in the expression of γ*CA1*, coding for a carbonic anhydrase involved in the mitochondrial respiratory chain, is measured in *swt-q* mutants in response to medium drought stress. In contrast, no significant change in the expression of both γ*CA1* and γ*CA2* is observed in WT (Table 2). Regarding chloroplast functioning, only slight changes in the expression of pPGI and PFK5 are observed in WT plants, while in swt-q mutant, a significant increase in the expression of *FRK3* is measured in response to drought (Table 2). In addition, both genotypes are significantly downregulating cytosolic isoforms of fructokinase (i.e., *FRK2* and *FRK7*) in response to drought, while a significant upregulation of cytosolic phosphofructokinases is measured only in WT plants in response to mild intensity drought (i.e., *PFK1* and *PFK7*) (Table 2). Finally, a significant upregulation of genes coding for enzymes involved in starch synthesis and degradation (i.e., *APL3*, *APL4* and/or *BAM1*) is measured in both genotypes in response to drought (Table 2). According to the slightly significant result of the GxE interaction this upregulation is further increased in *swt-q* mutant in response to medium-intensity drought stress (Table 2). This is accompanied by a significant upregulation of the expression of the gene coding for *MEX1*, a maltose transporter in both genotypes in response to drought. This upregulation is significantly more important in the *swt-q* mutant than the wild type, suggesting that in response to drought, the quadruple *sweet* mutant potentially synthesizes and degrades more starch and exports more maltose than WT plants (Table 2).

**Table 2.**
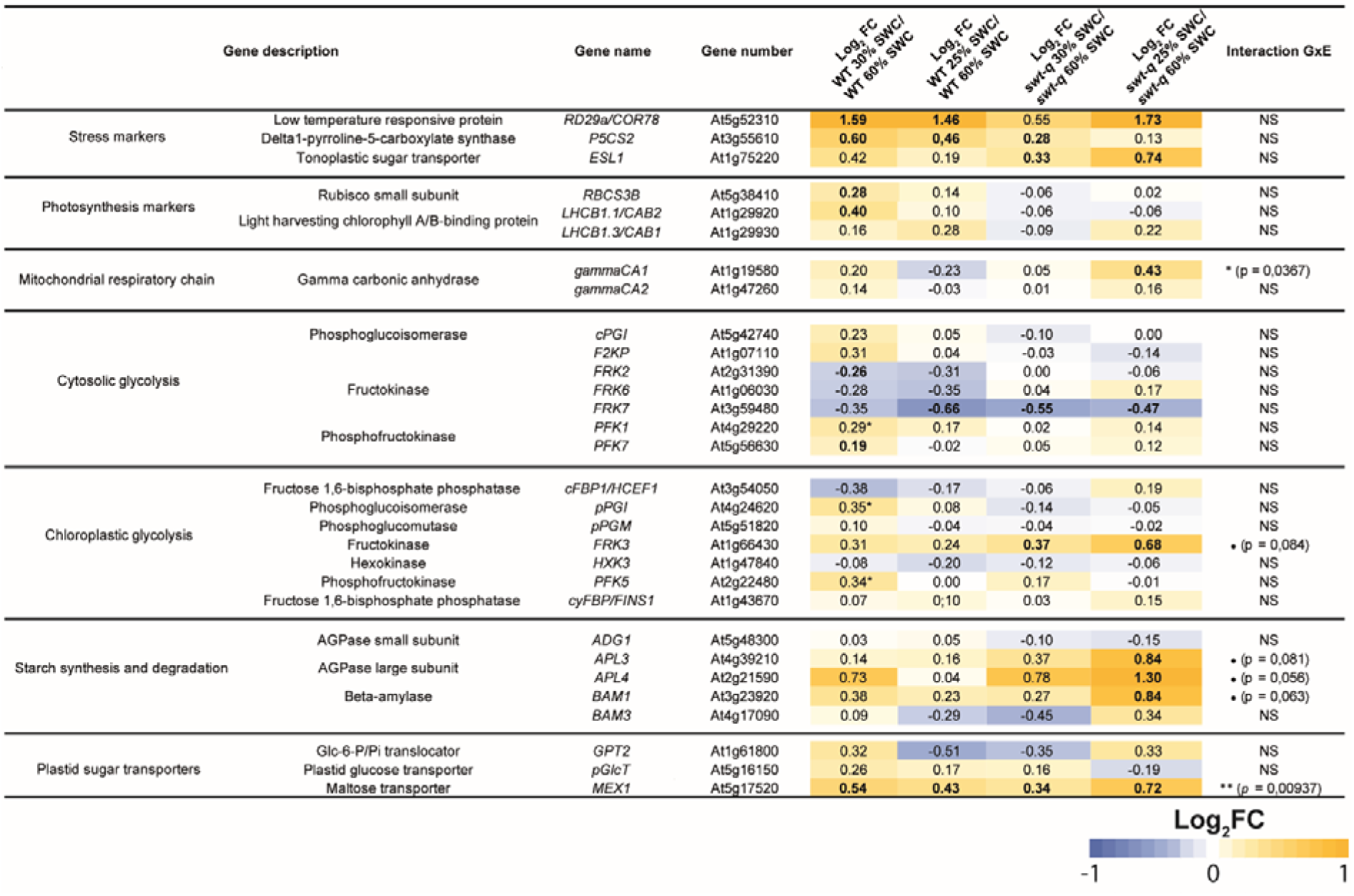
Log_2_-fold changes (FC) in the expression of genes involved in sugar homeostasis in the cytosol and the chloroplasts in the *swt-q* mutant and the wild type grown in mild-intensity (30% SWC) or medium-intensity (25% SWC) drought stress compared to normal watering conditions (60% SWC). Whole rosettes have been harvested for mRNA extraction and subsequent qPCR experiments. The RT-qPCR raw expression data are presented in Supplementary Table S9. Bold-scripted Log_2_FC indicate significant fold changes (*p* < 0.05), while stars indicate Log_2_FC values for which *P* is between 0.05 and 0.09 according to a double-sided *t*-test. The results of GxE interaction from the two-way ANOVA are also displayed (* *p* < 0.05, ** *p* < 0.01, • 0.05 < *p* < 0.09, NS: not significant).

## Discussion

The balance between soluble sugars and starch synthesis and utilization is crucial for plants to ensure appropriate growth under both normal and challenging environmental conditions (Dong and Beckles 2019). This equilibrium is achieved by the combined action of primary metabolism enzymes and intercellular and intracellular sugar transporters, among which transporters from the SWEET family. In this work, we further explore the phenotype of the sugar-accumulating *swt11swt12*, *swt16swt17,* and *swt-q* mutant lines (Chen et al. 2012; Gebauer et al. 2017; Aubry et al. 2022; Hoffmann et al. 2022) regarding their photosynthesis performance and response to drought stress since sugars can impact both (Goldschmidt and Huber 1992; Saddhe et al. 2021).

The seminal analysis of the *swt11swt12* mutant showed that SWEET11 and SWEET12 transporters are involved in phloem sugar loading in leaves, leading to an accumulation of sugars at the leaf level (Chen et al. 2012). In agreement, we show that the *swt11swt12* double mutant displays a significant accumulation of soluble sugars (i.e., sucrose, glucose, and fructose) and starch when grown in a short-day photoperiod. Since SWEET transporters transport sugars along the concentration gradient, it is important to get an idea in which subcellular compartment sugars are accumulating. Interestingly, our results strongly favor an accumulation of sugars in the cytosol of the *swt11swt12* double mutant (Fig. 6B). This hypothesis is sustained by the significant upregulation of the sugar-induced *GPT2, PFK5, ADG1, pPGI* and *pFBA3* together with the significant downregulation of the sugar-repressed *TPT* and *FBA7* (Table 1) (Weise et al. 2019; Khan et al. 2023 and Supplementary Table S10). Moreover, the expression pattern of *SWEET11* and *SWEE12* transcripts and proteins suggests that this accumulation likely takes place both in vascular parenchyma cells and rosette leaves’ mesophyll cells. Interestingly, this is accompanied by an increased loss of water per unit of biomass produced in the *swt11swt12* double mutant, as indicated by the lower WUE and WUEi. Unlike WUE, which in Arabidopsis has been shown to be positively linked to biomass accumulation, WUEi is more dependent on stomata opening/closure (e.g. anatomy, physiology, leaf biochemistry, and Calvin cycle activity) (Vialet-Chabrand et al. 2016; Bhaskara et al. 2022). In addition, the temperature of rosette leaves of the *swt11swt12* double mutant was lower than that of the wild type, suggesting an increased evapotranspiration (Supplementary Fig. S2). Altogether, these results support the hypothesis that stomata closure is impaired in the *swt11swt12* mutant compared to WT plants. Previous works point out that stomatal closure is sucrose-induced in most species, including Arabidopsis (Lawson et al. 2014; Kottapalli et al. 2018). It was also proposed that when the rate of sucrose synthesis exceeds the rate of sucrose loading into the phloem, the extra sucrose is carried toward the stomata by the transpiration stream to stimulate stomatal closure via hexokinase-dependent or independent mechanisms, preventing the loss of water and downregulating CO_2_ assimilation (Kelly et al. 2013; Lima et al. 2018). Here, we propose that in the *swt11swt12* mutant, the mechanisms allowing sucrose to reach the stomata via the transpiration stream are impaired, preventing the sucrose-induced stomatal closure from occurring and yielding lower WUE and WUEi (Fig. 6A-B). Consequently, since stomatal closure seems less efficient associated with a tendency for higher transpiration and stomatal conductance, it could account for the higher intracellular CO_2_ concentration (Ci) measured in the *swt11swt12* mutant (Fig. 2D). Finally, whether impaired function of quinone acceptors (Q_A_ and Q_B_), inactivation of the Mn_4_Ca cluster or the production of reactive oxygen species are responsible for the increased photodamage of the PSII observed in this mutant, needs to be further explored. Nonetheless, it could be hypothesized that, because of the negative regulation of the expression of Calvin cycle genes by sugar accumulation, a poor recycling of NADP^+^ and excessive electron transfer could take place, leading to ROS production and thus impacting PSII efficiency (Couée et al. 2006).

**Fig. 6.**
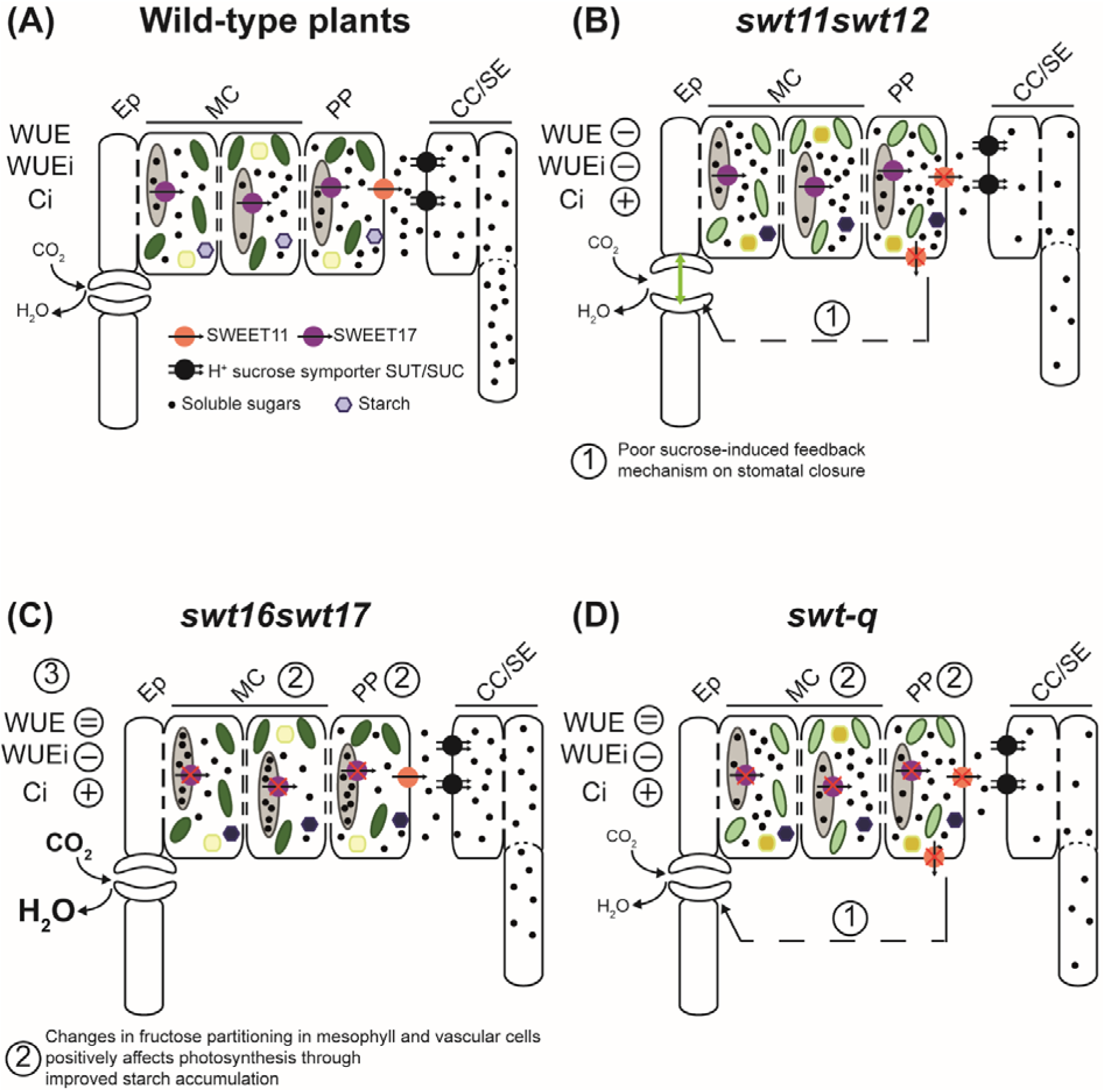
Proposed model for the role of SWEET11 and SWEET17 in photosynthesis. Synthesis scheme in wild-type plants (A), *swt11swt12* (B), *swt16swt17* (C), and *swt-q* (D) mutant lines. Within the mesophyll and phloem parenchyma cells, the vacuole is presented in light grey, the chloroplasts in green, the mitochondria in yellow, and a starch granule is depicted in purple. The light green double-headed arrow across the stomata in (B) refers to the potential increase in stomatal opening in the mutants. The differences in the color intensity of the chloroplasts, mitochondria, and starch granules represent changes in PSII efficiency, accumulation of organic acids, and starch accumulation, respectively. Finally, the size of the character (CO_2_, and H_2_O) refers to the increase compared to wild-type plants. To avoid overloading the figure, the organelles in the companion cell/sieve element complex are not presented. CC/SE: companion cells/sieve elements complex, Ep: epidermis, MCs: mesophyll cells, PP: phloem parenchyma cells.

In addition, in the *swt11swt12* mutant, the expression of several genes involved in cytosolic and plastidial glycolysis are also deregulated, which is associated to an altered TCA cycle functioning and to an increase of several organic acids content (e.g., malate, maleate, citrate, citramalate, 2-oxoglutarate, succinate, aconitate) as measured in the metabolomic analysis (Table 1 and Fig. 3). In complement to the ATP production during photosynthesis (through the chloroplastic electron transport chain, including the PSII), the TCA cycle generates the reducing equivalents NADH and FADH_2_ in the mitochondrial respiratory chain that fuel ATP synthesis by oxidative phosphorylation (Braun 2020). The carboxylic acid metabolism also provides carbon skeletons for nitrogen assimilation and aspartate biosynthesis. Interestingly, the content of several amino acids (e.g., aspartate, GABA, arginine, leucine, isoleucine, threonine, glycine) is slightly impaired in *swt11swt12* mutant (Fig. 3). Our results concur in proposing that in this mutant, the balance between ATP production (from both chloroplasts, due to the impaired PSII efficiency, and mitochondria) and N metabolism is modified likely at the expense of ATP since aspartate is slightly accumulating in the *swt11swt12* mutant (Fig. 3). Thus, a tight control of the cytosolic sugar content mediated by SWEET transporters is needed to maintain the energetic balance of the cell. Moreover, it could be proposed that the sugar exchanges between parenchyma cells and companion cells, mainly mediated by SWEET11, is also needed for the regulation of stomata functioning and the overall leaf photosynthesis. Thus, reinforcing the link between phloem loading, energetic status and their impact on stomatal closure.

So far, it has been suggested that, in the *swt16swt17* mutant, the vacuolar fructose content is likely enhanced at the expense of the cytosolic fructose content (Valifard et al. 2021; Aubry et al. 2022). Consistently, the expression of *pFBA1*, *pFBA2*, *FBA5*, *BAM1*, *pGlcT*, *PFK1,* and *F2KP* is downregulated in the *swt16swt17* mutant. Interestingly, the expression of these genes is upregulated in high cytosolic fructose content induced in fructose-incubated leaf discs (Supplementary Table S10 and Khan et al. 2023), supporting our hypothesis of low cytosolic fructose levels in *swt16swt17*. Consistenly the expression of the photosynthesis-related genes *LHCB1.1/CAB2* and *LHCB1.3/CAB1*, which are known to be repressed by high cytosolic sugar content (Supplementary Table S10), are upregulated in the *swt16swt17* mutant (Table 1). Thus, these results suggest that SWEET16 and SWEET17 act as fructose exporters in leaves when plants are grown in SD conditions, in a similar way as in roots of plants exposed to extended dark periods (Guo et al. 2014). On the contrary, while the expression of the plastidial isoforms of *ADG1*, *pFBA3*, *PFK5,* and *pPGI* is upregulated when cytosolic fructose content is high, their expression is also upregulated in the *swt16swt17* mutant (Table 1 and Supplementary Table S10). From these results, we postulate that fructose partitioning in the double mutant might be more complex than just increased vacuolar fructose sequestration and that the plastidial and/or cytosolic fructose content could also be impacted. Nonetheless, our results point out that vacuole-cytosol sugar exchanges play a crucial role in regulating the leaf’s gas exchanges (Fig. 6A and C). Indeed, in the *swt16swt17* mutant, several gas exchanges-related parameters are improved compared to wild-type plants (i.e., net CO_2_ assimilation, transpiration rate, stomatal conductance, and Ci) (Fig. 2). Consequently, we propose that the modified fructose partitioning within the mesophyll cells and/or the vascular system positively impact the plant photosynthesis (Fig. 6C). One possible explanation for this could reside in the higher starch synthesis measured in the *swt16swt17* mutant (Table 1 and Fig. S4). Indeed, the expression of *pFBA1-3, pPGI,* and *ADG1* genes, involved in starch biosynthesis (Lu et al. 2012; Preiser et al. 2020) are significantly upregulated (Table 1). Moreover, despite the slight upregulation of *BAM1* expression and that of *pGlcT* shown to export glucose after starch degradation (Cho et al. 2011), increased starch content is measured in the *swt16swt17* mutant (Table 1 and Supplementary Fig. S4). Because high photosynthesis correlates with the up-regulation of starch metabolism genes (MacNeill et al. 2017) and that fructose and starch synthesis are linked (e.g., FINS/FBP, F2KP) (Cho and Yoo 2011; McCormick and Kruger 2015), the increased starch accumulation in the *swt16swt17* mutant could also account for the improved photosynthesis (Fig. 6C).

It is well known that accumulation of sugars, acting as osmolytes or ROS scavengers, is a common response to protect plants subjected to abiotic stresses (Keunen et al. 2013; Saddhe et al. 2021). However, it has also been shown that, in response to drought stress, soluble sugars are not the main contributors to osmotic adjustment unlike potassium, organic acids (i.e. fumarate) or amino acids (i.e, proline) (Hummel et al. 2010). In Arabidopsis *swt11swt12* and *swt17* mutants, which accumulate higher levels of sugars, have been shown to be more sensitive to drought stress (Valifard et al. 2021; Chen et al. 2022). On the other hand, the overexpression of the apple tonoplastic MdSWEET17 in tomato leads to an accumulation of sugars (likely through increased export of sugars to the cytosol), which improves the tomato’s response to drought (Lu et al. 2019). These works suggest that the response to drought is not only linked to an overall increase of sugar content in leaves but could also depend on the sugar partitioning between vacuole and cytosol. When the *swt-q* mutant is grown in normal watering conditions, its fructose content is higher than that of the double mutants, suggesting an additive phenotype with regard to fructose accumulation (Fig. 3). Among the 7 genes significantly deregulated in the *swt-q* mutant, *ADG1*, *PFK5*, *GPT2* and *MEX1,* induced by high cytosolic fructose/glucose content (Supplementary Table S10), are up-regulated (Table 1). These results suggest that in the quadruple mutant, and in the *swt11swt12* double mutant, higher sugar accumulation is likely taking place in the cytosol rather than in the vacuole. We propose that the impaired sugar partitioning in the *swt-q* mutant accounts for an increased sensitivity to drought even when imposed at a mild intensity (30 % SWC) (Fig. 4). This is in agreement with previous work showing that *swt11swt12* displays a reduced shoot fresh weight in response to drought stress when grown in soil (Chen et al. 2022 and Supplementary Fig. S5). On the contrary an increased fructose trapping in the vacuole, as proposed for the *swt16swt17* double mutant, slightly improves the plant resistance to drought (Supplementary Fig. S5). This result was surprising since we previously showed that the *swt17* single mutant was more sensitive to drought than the wild type (Valifard et al. 2021, 2024). This discrepancy could be due to compensatory mechanisms, yet to be identified, which could improve the plant response to drought. Nonetheless, these results support the hypothesis that tight control of the balance between cytosolic and vacuolar sugar content is indispensable for the plant’s response to drought.

Besides impacting the plant morphology, drought stress affects plant physiology, impacting photosynthesis as a consequence of stomata closure (Kaur et al. 2021). Moreover, a higher intrinsic water use efficiency (WUEi) linked to improved root growth is often associated with drought tolerance (Gago et al. 2014). Consequently, the reduced root growth observed in both *sw11swt12* and *swt17* mutant lines was proposed to be responsible for the reduced drought resistance. However, the primary root length is not affected in the *swt16swt17* double mutant when grown *in vitro* (Guo et al. 2014) while its response to drought is slightly improved compared to WT. In the *swt-q* mutant, the reduction of the WUEi could also explain the increased sensitivity to drought, but a more detailed analysis of the root system development needs to be performed to get an integrative view of the SWEET-mediated response to drought. Despite the initial impaired photosynthesis performance (Fig. 6D), the photosynthesis does not seem to be further affected by drought in *swt-q* mutant compared to the WT, since no change in the expression of photosynthesis marker genes is measured (Table 2). Nonetheless, the *swt-q* mutant accumulates significantly more sucrose and glucose than the WT when exposed to drought (Fig. 5). In addition, the expression of genes involved in starch synthesis and degradation (i.e., *APL3*, *APL4*, *BAM1* and *MEX1*) is further upregulated in *swt-q* mutant compared to the wild type when subjected to drought (Table 2). Interestingly, the stable expression of the sugar-repressed *LHCB1.3*/*CAB1* and sugar-induced *GPT2,* under drought compared to WT, suggests that the cytosolic sugar level in *swt-q* mutant is not further modified in response to drought. Interestingly, the expression of *ERDL3.07/ESL1*, known to import sugars in the vacuole in response to drought stress (Yamada et al. 2010; Slawinski et al. 2021), is also significantly upregulated in *swt-q*, thus limiting the adverse effect of drought on photosynthesis inhibition.

## Conclusion

It has long been proposed that changes in sugar accumulation impact plant photosynthesis performance. Nonetheless, the molecular actors involved in such a phenomenon are not well characterized. In this work, we propose that a higher cytosolic sugar accumulation in the mesophyll and phloem parenchyma cells takes place in the *swt11swt12* mutant line, leading to an altered sucrose-induced feedback mechanism on stomatal closure. On the other hand, we propose that the increased fructose sequestration in the vacuole of the *swt16swt17* double mutant positively impacts the plant photosynthesis performance most probably by uplifting the photosynthesis inhibition by its end-products. Furthermore, this work proposed that the impaired sugar partitioning observed in the *swt-q* quadruple mutant is associated with an enhanced sensitivity upon drought.

## Author Contributions

Conceptualization, R.L.H.; investigation, E.A., G.C. and R.L.H.; methodology, E.A., G.C., E.G., and R.L.H.; visualization, R.L.H.; writing—original draft, R.L.H.; writing—review and editing, S.D. and R.L.H.

## Acknowledgments

We thank Olivier Loudet (head of the *Phenoscope* platform) for his help in validating the results of the *Phenoscope* experiments, and Catherine Bellini for critically reading the manuscript.

## Data availability

The data supporting the findings of this study are available from the corresponding author, Rozenn Le Hir, upon request.

## Supplementary data

Fig. S1. Rosette growth of *sweet* mutant lines grown under short-days photoperiod.

Fig. S2. Maximum rosette leaves temperature is impaired in *swt11swt12* mutant line.

Fig. S3. Expression of SWEET transporters in the rosette leaves grown under short-days photoperiod.

Fig. S4. Starch accumulates in rosette leaves of *sweet* mutant lines.

Fig. S5. Changes in projected rosette area of the different mutant lines in response to drought.

Fig. S6. Proline synthesis is upregulated in response to drought stress.

Fig. S7. Metabolites specifically accumulating or decreasing in response to different drought stress in wild type or *swt-q* mutant.

Table S1. Primers used for quantifying genes by RT-qPCR.

Table S2. Metabolites significantly different in the different *sweet* mutant lines compared to wild-type plants grown in short-days photoperiod under normal watering conditions.

Table S3. Raw data of the *Phenoscope* experiment.

Table S4. Metabolites significantly different in wild-type plants grown in non-limiting conditions (60% SWC) or after the application of a long-term mild intensity stress (30% SWC).

Table S5. Metabolites significantly different in wild-type plants grown in non-limiting conditions (60% SWC) or after the application of a long-term medium intensity stress (25 % SWC).

Table S6. Metabolites significantly different in *swt-q* plants grown in non-limiting conditions (60% SWC) or after the application of a long-term mild intensity stress (30 % SWC).

Table S7. Metabolites significantly different in *swt-q* plants grown in non-limiting conditions (60% SWC) or after the application of a long-term medium intensity stress (25 % SWC).

Table S8. Metabolites significantly different between both genotypes and both drought stress conditions.

Table S9. RT-qPCR raw expression data in normal growth condition.

Table S10. Sugar response of genes significantly deregulated in *sweet* mutant lines.

