## Supplementary-figures for "Changes in SWEET-mediated sugar partitioning affect photosynthesis performance and plant response to drought"

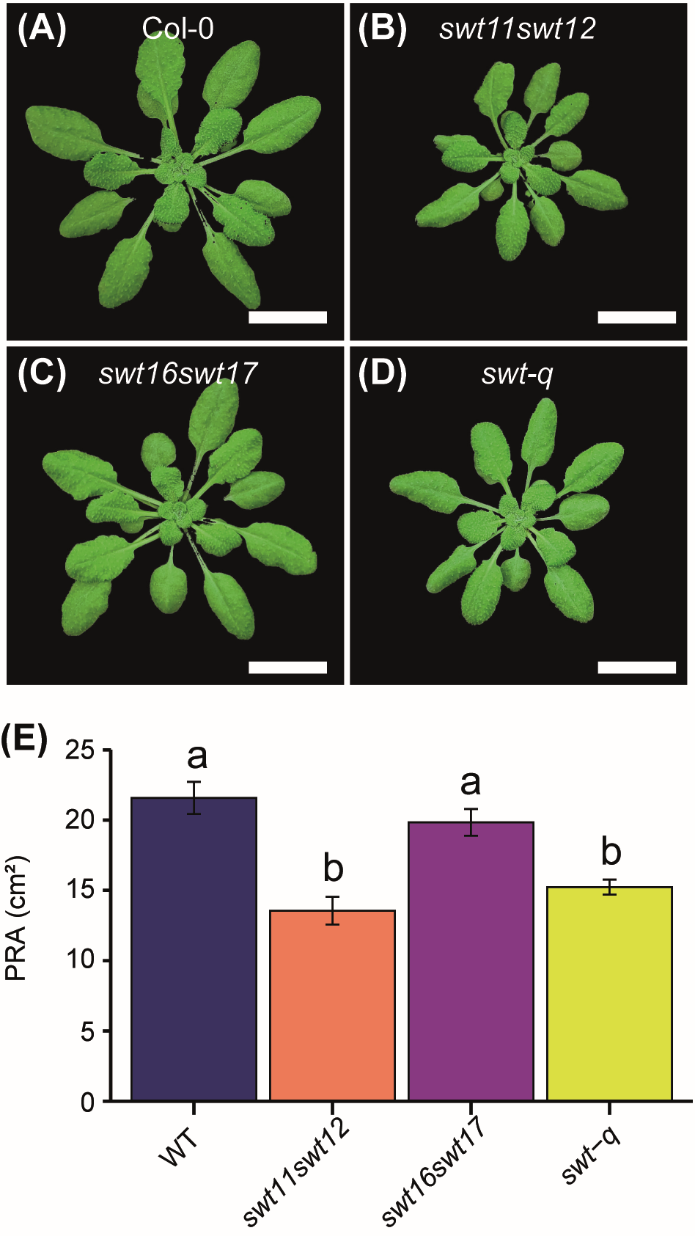

**Supplementary Fig. S1. Rosette growth of *sweet* mutants grown under short-day photoperiod. (**A-D) Representative picture of a wild-type plant (A), *swt11swt12* (B), *swt16swt17* (C), and *swt-q* mutants (D) grown under short-day conditions for 43 days. Scale bar = 2 cm. (E) Barplots showing the projected rosette area (PRA). Values are means ± SE (n ≥ 23 plants from two independent cultures). Significant differences were calculated using a one-way ANOVA combined with Tukey’s comparison post hoc test.

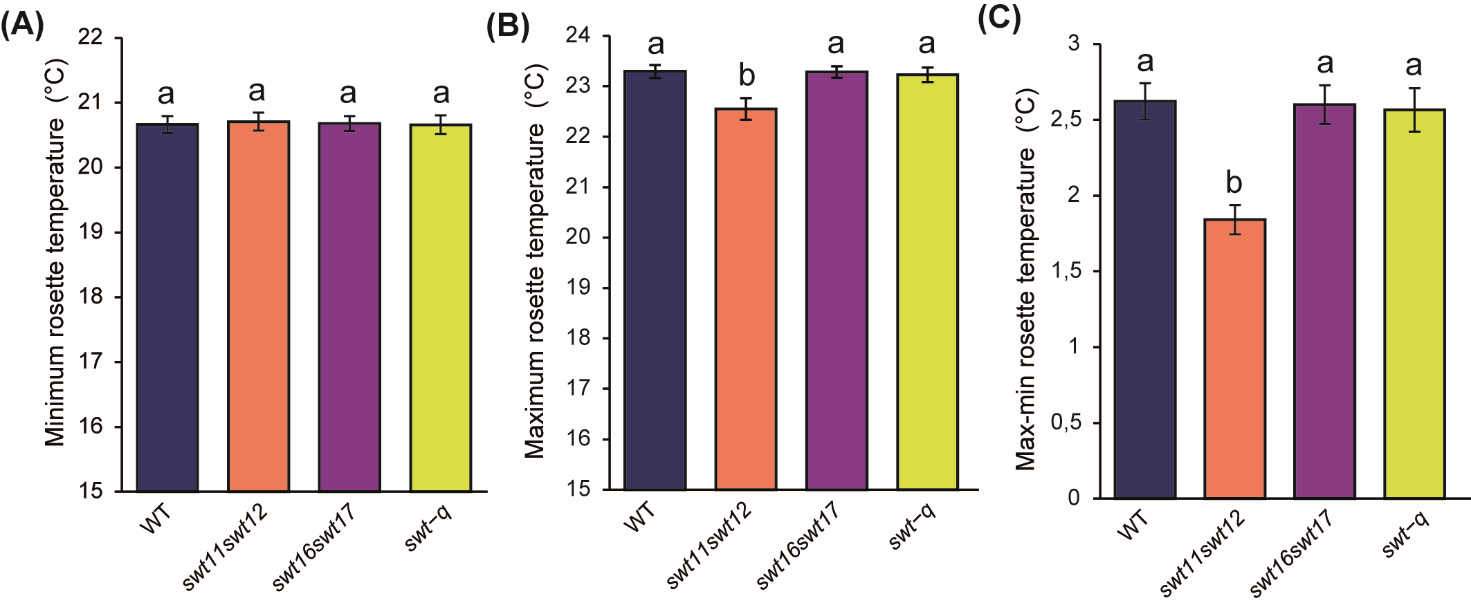

**Supplementary Fig. S2. Maximum temperature of rosette leaves is impaired in *swt11swt12* mutant.** Barplots showing the minimum whole rosette temperature (A), the maximum whole rosette temperature (B), and the difference between the whole rosette maximum and minimum temperature (C) of wild type, *swt11swt12*, *swt16swt17,* and *swt-q* mutants grown in short-day photoperiod. Means ± SE are shown (n ≥ 23 plants per genotype from 2 independent experiments). A one-way ANOVA combined with Tukey’s comparison post hoc test was performed. The values marked with the same letter were not significantly different from each other, whereas different letters indicate significant differences (*p* < 0.05).

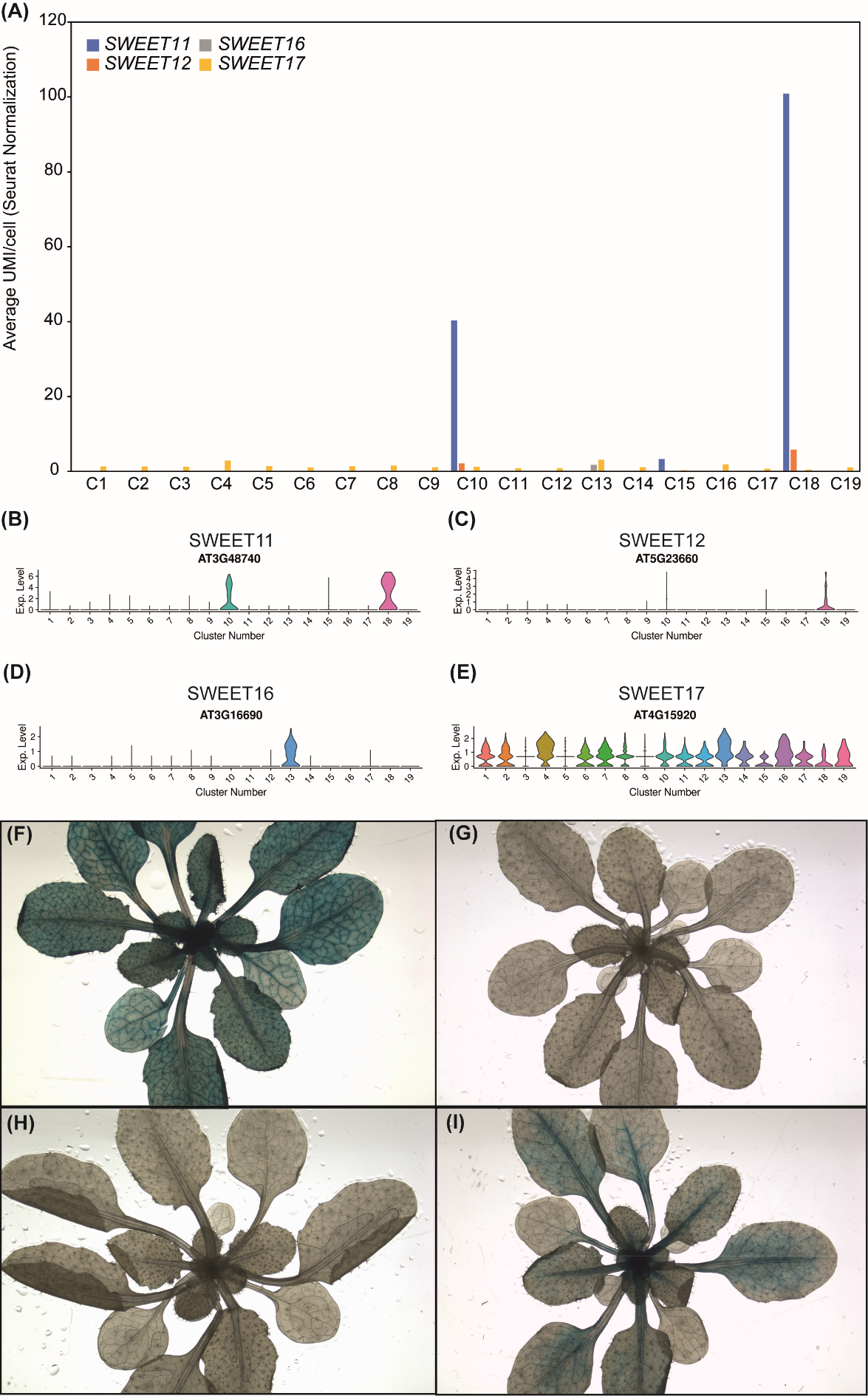

**Supplementary Fig. S3. Expression of SWEET transporters in the rosette leaves grown under short-day photoperiod.** (A) Barplot showing the average unique transcripts per cell (UMI stands for Unique Molecular Identifiers) after Seurat normalization which reflects the expression level of each gene. Single-cell leaf atlas has been performed on Arabidopsis rosette leaves grown in short-days photoperiod. Clusters 1, 2, 3, 5, 6, 7, 8, 9, 11, 12 and 14 refer to mesophyll cells, cluster 4 to bundle sheath and xylem, cluster 10 to phloem parenchyma and procambium, cluster 13 to epidermis, cluster 18 to phloem parenchyma and xylem, cluster 16 to guard cells, cluster 15 to companion cells, cluster 17 to hydathode and cluster 19 to unassigned cluster. For more details see (Kim et al. 2021) and the Plant scRNA-seq database website (<https://www.zmbp-resources.uni-tuebingen.de/timmermans/plant-single-cell-browser-leaf-atlas/>).

(B-E) Violin plots depicting the distribution of expression levels for *SWEET11* (B), *SWEET12* (C), *SWEET16* (D), and *SWEET17* (E).

(F-I) Histochemical analysis of GUS activity in lines expressing SWEET-GUS fusion proteins driven by their native promoter in rosettes leaves of plants grown in short-days photoperiod, namely pSWEET11:SWEET11-GUS (F), pSWEET12:SWEET12-GUS (G), pSWEET16:SWEET16-GUS (H) and pSWEET17:SWEET17-GUS (I).

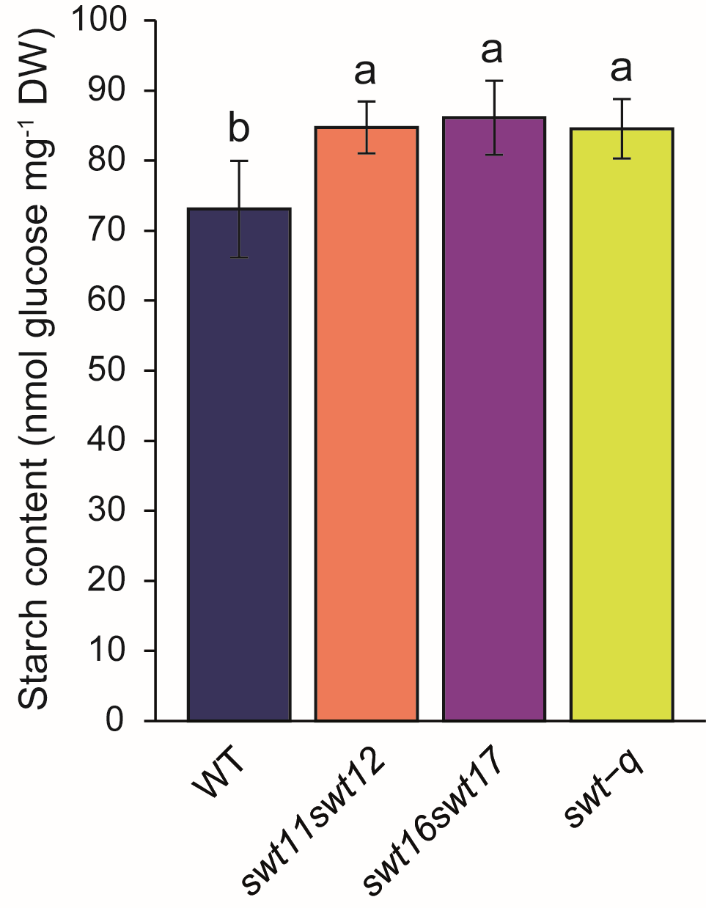

**Supplementary Fig. S4. Starch accumulates in rosette leaves of *sweet* mutants.** Barplot shows the starch content in wild type and *sweet* mutant rosette leaves grown in a short-day photoperiod for 43 days. Means ± SD are shown (n = 6 plants per genotype). A one-way ANOVA combined with Tukey’s comparison post hoc test was performed. The values marked with the same letter were not significantly different from each other, whereas different letters indicate significant differences (*p* < 0.05).

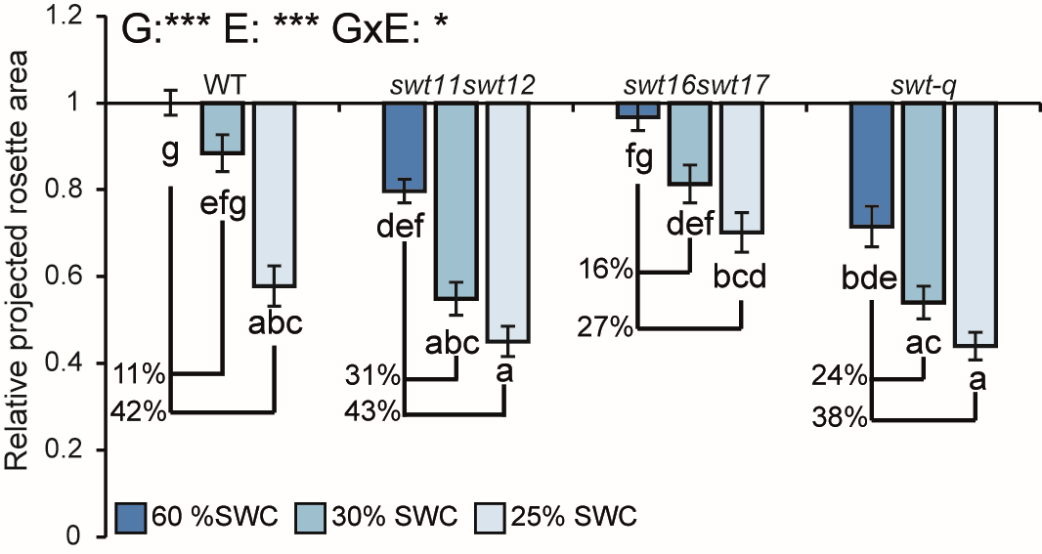

**Supplementary Fig. S5. Changes in the wild type and the different mutants projected rosette area in response to drought.** Barplot showing the fold-changes in the projected rosette area of the wild type, *swt11swt12*, *swt16swt17,* and *swt-q* mutants grown in short-day photoperiod under different watering regime: 60% SWC (normal watering conditions), 30 % SWC (mild stress intensity) or 25% SWC (medium stress intensity). The fold changes were determined using the values that were normalized to the mean of wild-type plants grown at 60% SWC. A two-way ANOVA combined with the Tukey’s comparison post hoc test was performed. The values marked with the same letter were not significantly different from each other, whereas different letters indicate significant differences (*p* < 0.05, *n* ≥ 18 per genotype and condition from 2 independent experiments).

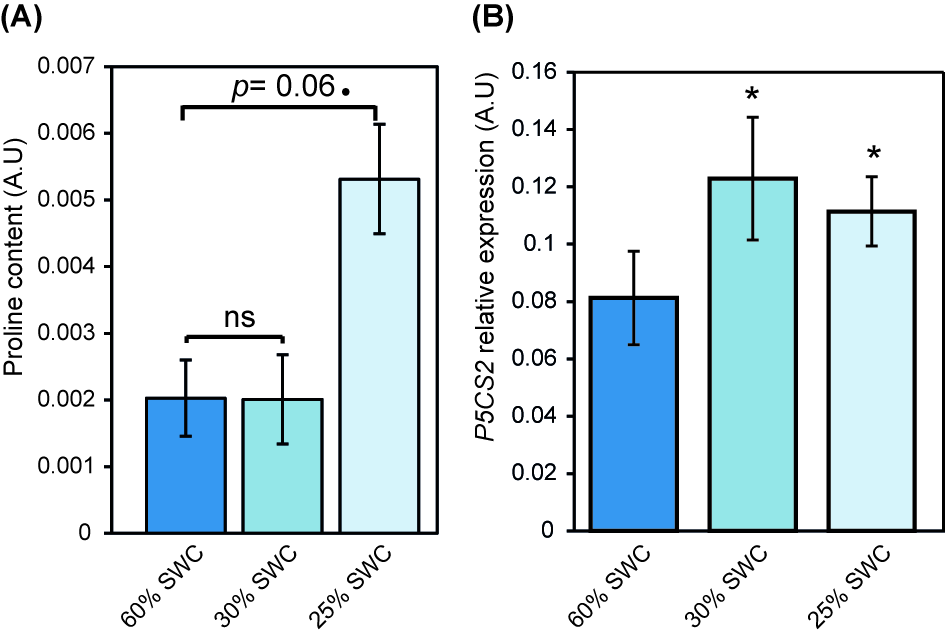

**Supplementary Fig. S6. Proline synthesis is upregulated in response to drought stress.** Barplots showing the proline content (A) and the expression of the *P5CS2* gene involved in prolonged biosynthesis (B) in wild-type plants grown under different water regimes: 60% SWC (normal watering conditions), 30 % SWC (mild stress intensity) or 25% SWC (medium stress intensity). The values are means ± SD (*n*=8 or 4 in (A) and (B) respectively) The stars represent the result of a student t*–*test to compare wild type grown at 60 % SWC and 30 % SWC or 25 % SWC (* *p* < 0.05, • 0.05 < *p* < 0.09, ns: not significant).

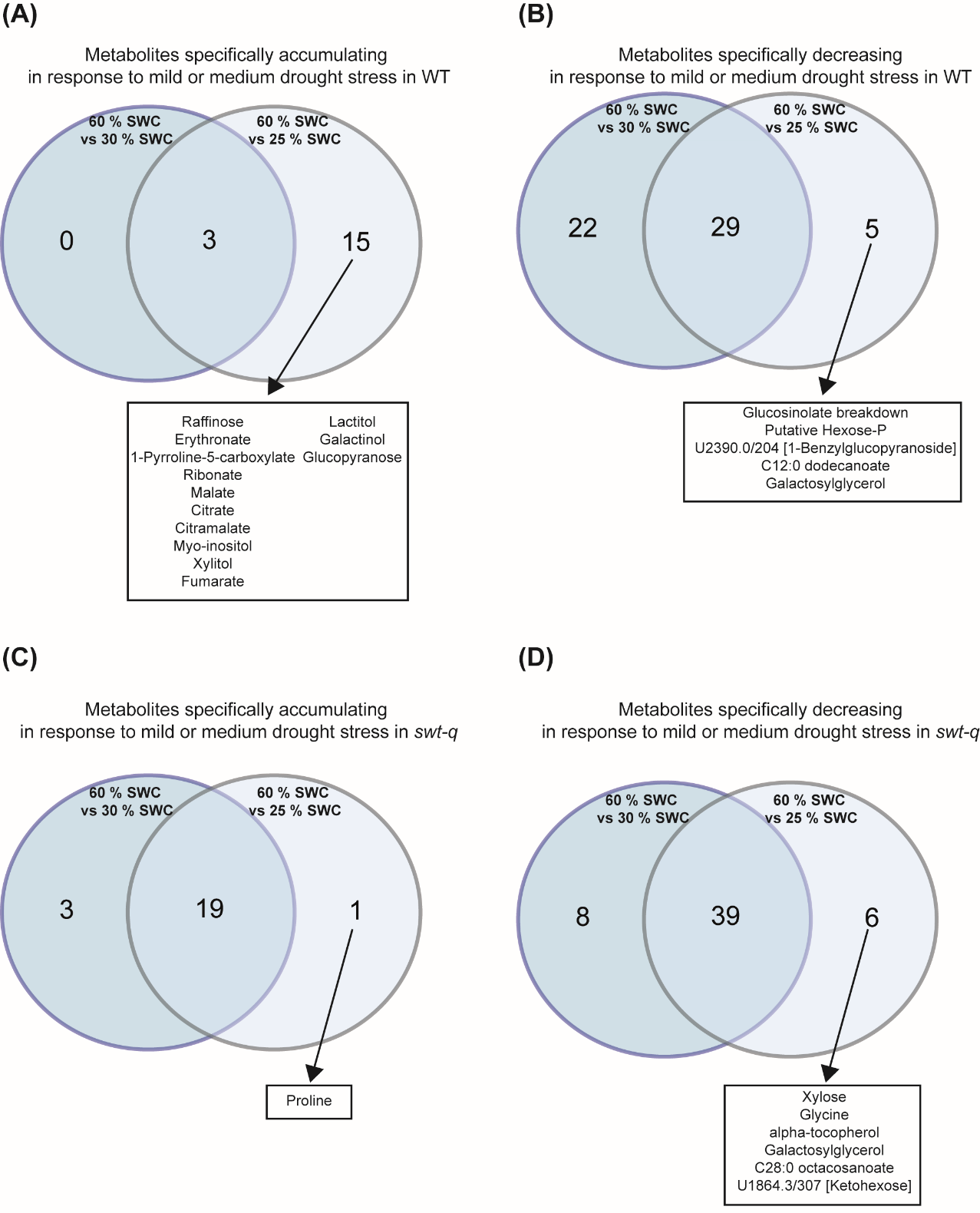

**Supplementary Fig. S7. Metabolites specifically accumulating or decreasing in response to different drought stress in the wild type or the *swt-q* mutant.** Venn diagram showing the metabolites specifically and significantly accumulating (A and C) or decreasing (B and D) in wild-type plants (A and B) or in *swt-q* mutant (C and D) subjected to mild intensity or medium intensity drought stress. The statistical analysis is presented in Supplemental Tables S2-5.
